# Suppression of YAP/TAZ molecular targets by epigenetic editing interferes with GBM growth and invasiveness

**DOI:** 10.64898/2026.03.12.711340

**Authors:** Eva Baronchelli, Silvia Ferretti, Michal Kubacki, Antonella Covino, Valerio Benedetti, Edoardo Bellini, Serena Gea Giannelli, Mirko Luoni, Elisa Ventura, Federica Banfi, Gaia Colasante, Federica Ungaro, Vania Broccoli, Alessandro Sessa

**Affiliations:** Stem Cells and Neurogenesis Unit, Division of Neuroscience, IRCCS San Raffaele Scientific Institute, Milan, 20132, Italy; Neuroepigenetics Unit, Division of Neuroscience, IRCCS San Raffaele Scientific Institute, Milan, 20132, Italy; Experimental Gastroenterology Unit, Department of Gastroenterology and Digestive Endoscopy and Division of Immunology, Transplantation and Infectious Disease, IRCCS San Raffaele Scientific Institute, Milano, 20132, Italy; CNR, Institute of Neuroscience, Milan, 20129, Italy

## Abstract

Glioblastoma (GBM) is a highly aggressive brain tumor characterized by extensive heterogeneity, diffuse invasion, and recurrence despite multimodal therapy. Aberrant transcriptional programs driven by oncogenic signaling pathways sustain GBM growth, stemness, and therapy resistance, yet targeting individual molecular nodes has yielded limited clinical benefit. Here, we introduce a transcriptional rewiring strategy based on an engineered epigenetic silencer factor (ESF) targeting the YAP/TAZ–TEAD axis. We developed a TEAD1 Epigenetic Silencer (TES) by fusing the DNA-binding domain of TEAD1 to repressive epigenetic modules. TES selectively binds TEAD genomic targets and imposes stable transcriptional repression of YAP/TAZ-dependent gene programs through chromatin remodeling and DNA methylation. Genome-wide analyses revealed that TES preserves TEAD1 DNA-binding specificity while converting an oncogenic transcriptional platform into a repressive state. Functionally, TES impaired proliferation, induced cell death, and reduced migratory and invasive properties in glioma cell lines and patient-derived cancer stem–like cells. *In vivo*, TES significantly reduced tumor growth in orthotopic GBM xenograft models and enhanced the therapeutic efficacy of temozolomide. Importantly, TES was well tolerated by normal neural cells *in vitro* and in the adult mouse brain *in vivo*. These findings establish TES as a proof-of-concept epigenetic therapy to durably suppress oncogenic transcriptional networks in GBM.

## INTRODUCTION

Glioblastoma (GBM) is the most common primary brain tumor in adults, accounting for approximately 50% of all malignant central nervous system neoplasms^1^. GBM remains one of the most lethal human cancers, as patient survival after diagnosis is still dramatically low despite aggressive treatment. Recurrence represents the major cause of therapeutic failure^2^. The current standard of care consists of maximal surgical resection followed by radiotherapy and temozolomide (TMZ) chemotherapy^3^. Recently approved therapies have only marginally extended patient survival, likely due to the highly infiltrative nature of the tumor and the limited efficacy of systemically delivered agents imposed by the blood-brain barrier (BBB)^4,5^.

Rapid advances in biotechnology have led to a broad range of experimental strategies aimed at eliminating or neutralizing tumor cells^5^. These include: (i) chemical or biological inhibitors targeting EGFR, VEGF, receptor tyrosine kinases (RTKs), or the PI3K pathway; (ii) immunotherapies, such as immune checkpoint inhibitors and CAR-T cell approaches; (iii) oncolytic viruses; and (iv) gene therapies. However, the limited translation of these approaches from experimental models to the clinic highlights the need for alternative strategies that may be combined with existing treatments.

GBM cancer stem–like cells (CSCs) have been implicated in disease aggressiveness and recurrence^6,7^. CSCs exhibit neural stem cell-like properties, including self-renewal, and display intrinsic or acquired resistance to radio- and chemo-therapy. They also help establish an immunosuppressive microenvironment that is beneficial for tumor growth^8–10^. Their infiltrative behavior enables them to escape surgical resection, as tumor cells often extend beyond the resection margin, contributing to relapses^11^.

Multiple pathways have been identified as central to CSC malignancy. Several transcriptional programs and associated transcription factors (TFs) regulate proliferation, migration, and invasion in cultured CSCs, patient-derived xenografts (PDX), and patient samples^12,13^. Chromatin-remodeling-driven plasticity has been linked to the regulation of these pathways and to aberrant CSC fate and behavior^14^. Importantly, CSCs display marked molecular heterogeneity, reflecting the intrinsic complexity of GBM, patient-specific features, and treatment-induced evolutionary dynamics^10,15,16^. This complexity likely explains why targeting individual genes yields limited therapeutic benefit, whereas approaches that modulate entire transcriptional programs or pathways tend to produce more robust effects.

In this context, we recently introduced an engineered epigenetic silencer factor (ESF) designed to repress the entire transcriptional output of SOX2, a TF with essential roles in GBM biology^17^. Such synthetic “epieditors”^18–21^ consist of epigenetic effector domains derived from transcriptional repressors fused to the SOX2 DNA-binding domain, producing the SOX2 Epigenetic Silencer (SES). SES induces a durable repressive chromatin state at SOX2 target *loci*, leading to reduced proliferation and cell death in GBM cells both *in vitro* and in PDX models^17^. This strategy was conceived as a cell-autonomous, cell therapy-like approach to eliminate or neutralize cancer cells, including CSCs, when used in combination with existing treatments to limit recurrence. Expanding the ESF toolkit could help overcome therapy resistance and improve therapeutic efficacy.

The Hippo signaling pathway is an evolutionarily conserved regulator of organ size, tissue homeostasis, and regeneration, acting through the control of stemness, proliferation, migration, and apoptosis^22,23^. In mammals, the pathway comprises tumor suppressors such as NF2, MST1/2, SAV1, and LATS1/2, which form a kinase cascade culminating in the phosphorylation and inactivation of the oncogenic coactivators YAP and its paralog TAZ. YAP/TAZ shuttle between cytoplasm and nucleus, where, in association with TEA domain (TEAD) transcription factors, they activate target gene transcription^23^. Hyperactivation of YAP/TAZ is widespread across cancers, where it promotes tumor initiation, progression, and metastasis^23–25^.

Here, we present a novel ESF based on the DNA-binding activity of TEAD1 and equipped with two repressive epigenetic modules: (i) the Kruppel-associated box (KRAB) domain, which promotes repressive histone modifications (e.g., increased H3K9me3 and removal of H3K4ac)^26^, and (ii) the catalytic domain of the DNA methyltransferase DNMT3A together with its cofactor DNMT3L, which induces DNA methylation^27^. We demonstrate that this TEAD1 Epigenetic Silencer (TES) binds DNA, represses YAP/TAZ target genes through DNA methylation, and reduces tumor burden in GBM models both *in vitro* and *in vivo*. Moreover, TES impairs both the proliferative and invasive capabilities of GBM cells, suggesting its potential to limit both tumor regrowth and dissemination during recurrence.

## RESULTS

### Generation of a TEAD1 epigenetic silencer

Following the strategy previously adopted for the generation of the SOX2 Epigenetic Silencer (SES)^17^, we engineered a TEAD1 Epigenetic Silencer (TES) by fusing the Kruppel-associated box (KRAB) domain, the catalytic domain of DNMT3A together with its cofactor DNMT3L, and the N-terminal 166 amino acids of TEAD1. This design excludes the C-terminal region of TEAD1, which mediates YAP/TAZ binding and transcriptional activation^28^ (**Figure 1A**). Computational modeling of TES was performed using the I-TASSER platform, based on homology modeling algorithms^29,30^ (**Table 1**). The top-ranked predicted structure was subsequently used to assess DNA-binding capability through molecular docking simulations using the HADDOCK software^31^. *In silico* analyses predicted a stable protein folding and preservation of the TEAD1 DNA-binding domain conformation despite the fusion of epigenetic effector domains, indicating maintained binding to the DNA double helix (**Figure 1B**).

**Figure 1.**
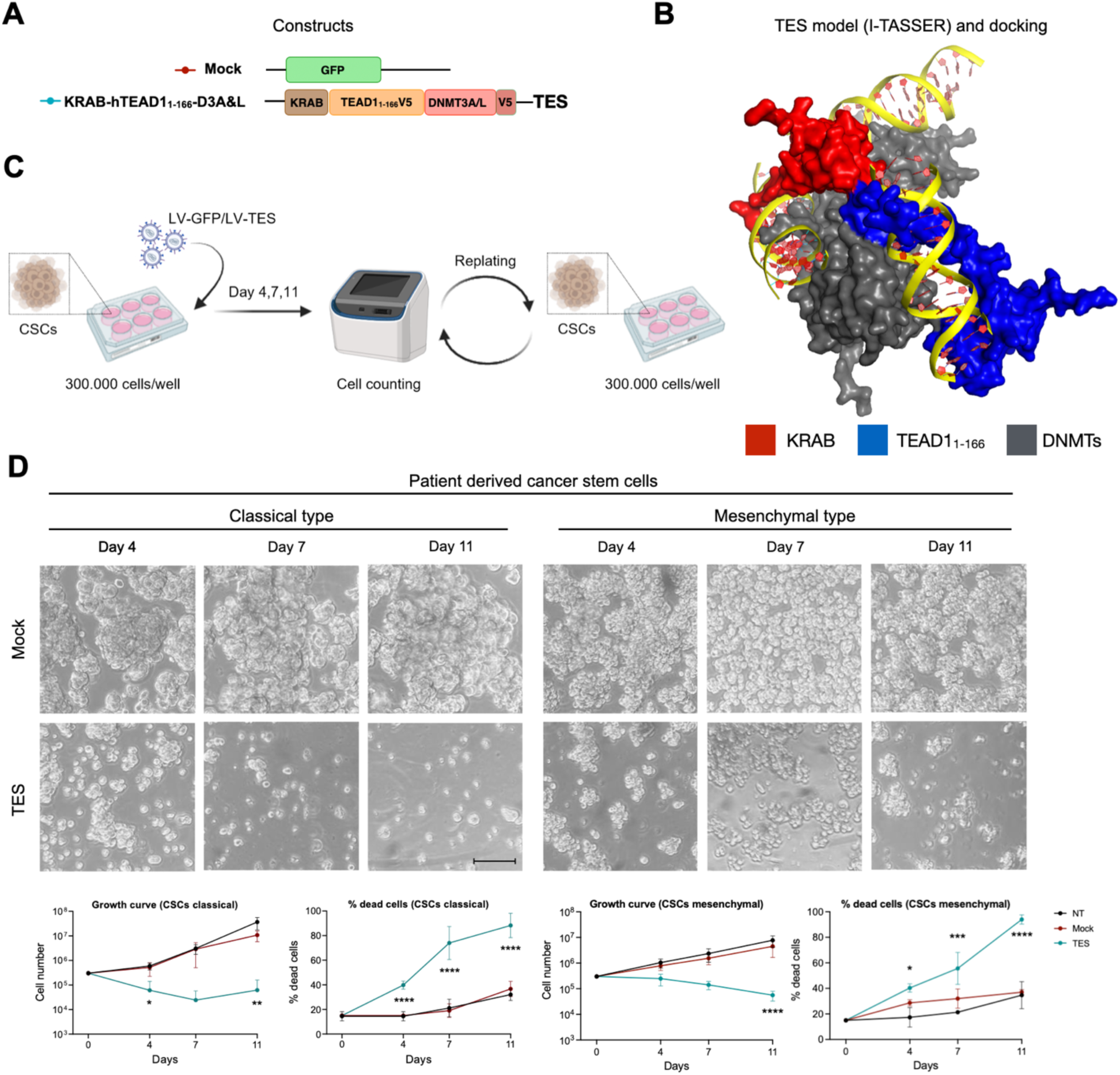
Generation and *in vitro* efficacy of the TEAD1 epigenetic silencer. (A) Schematic representation of the TES construct generated by fusing the N-terminal region of the human TEAD1 TF with the KRAB repressor domain at its N-terminus, and with DNMT3A and DNMT3L catalytic domains at its C-terminus. A V5 epitope tag was added at the C-terminus of the transgene. A GFP-expressing construct was used as mock control. (B) Docking simulation between the TES protein model and 19 DNA nucleotides. Here the best pose, filtered by 𝚫G of binding, is shown. This analysis supports the possibility for TES to bind DNA (shown in yellow). The human TEAD1 domain is highlighted in blue, whereas the KRAB domain and DNMTs catalytic domains are shown in red and grey, respectively. (C) Experimental scheme of the infection protocol used in patient-derived GBM CSCs. (D) *(Left)* Representative bright-field images, growth curve, and percentage of dead cells (trypan blue automatic counting) of patient-derived CSCs of classical GBM subtype infected with GFP (mock, in red), TES (in blue), or not treated cells (NT, in black), indicating strong capability of TES in inhibiting cCSCs growth and proliferation. Growth curve: Mock vs TES Day 4 \**P*=0.0196; Mock vs TES Day 7 *P*=0.0554; Mock vs TES Day 11 \*\**P*=0.0057. Dead cells: Mock vs TES \*\*\*\**P*<0.0001. *(Right)* Representative bright-field images, growth curve, and percentage of dead cells (trypan blue automatic counting) of patient-derived CSCs of mesenchymal GBM subtype infected with GFP (mock, in red), TES (in blue), or not treated cells (NT, in black), indicating strong capability of TES in inhibiting mCSCs growth and proliferation. Growth curve: Mock vs TES Day 4 *P*=0.7718; Mock vs TES Day 7 *P*=0.2092; Mock vs TES Day 11 \*\*\*\**P*<0.0001. Dead cells: Mock vs TES Day 4 \**P*=0.0429; Mock vs TES Day 7 \*\*\**P*=0.0001; Mock vs TES Day 11 \*\*\*\**P*<0.0001. Data are expressed as means ± SEM. Statistical analysis was performed using two-way ANOVA followed by Dunnett’s multiple comparisons test. *n* = 6. Scale bar: 100 μm.

Lentiviral (LV) transduction of TES into the SNB19 glioma cell line *in vitro* (**Figure S1A**) resulted in robust expression of the exogenous factor, as detected by the V5 epitope tag included in the construct (**Figure S1B,C**). TES expression significantly impaired SNB19 cell growth (**Figure S1D,E**) and similarly reduced proliferation in the U87 glioma cell line (**Figure S1F,G**). Proliferative reduction was blunted when the mutation R85K^32^ was inserted in TES to inhibit DNA binding (TES mut), thus confirming that TES anti-tumor activity is driven by its DNA-binding capacity (**Figure S1D,E**). We next extended this analysis to patient-derived GBM cancer stem-like cells (CSCs) representing both classical and mesenchymal molecular subtypes, which more faithfully preserve features of the primary tumors (**Figure 1C**). Efficient LV-mediated transduction was achieved in all CSC cultures using either LV-TES or LV-GFP as a control (**Figure S2**). TES expression resulted in a marked reduction in proliferative capacity accompanied by extensive cell death across both CSC subtypes (**Figure 1D** and **Figure S2**).

These results demonstrate that an ESF based on the TEAD1 transcription factor effectively suppresses glioma cell growth and induces cell death *in vitro*.

### TES molecular function

To elucidate the molecular program underlying TES biological activity, we analyzed the global transcriptional profile of SNB19 cells 48 hours after LV administration. TES expression induced widespread transcriptional alterations, with several thousand genes significantly up-or downregulated compared to control cells (**Figure 2A, Figure S3A,B,** and **Table S2**). Gene Ontology (GO) and Gene Set Enrichment Analysis (GSEA) revealed that the downregulated gene set was markedly more coherent, with numerous significantly enriched terms, whereas upregulated genes showed limited functional enrichment (**Figure 2B**, and **Figure S3C,D**). This pattern is consistent with the expected role of TES as a transcriptional repressor. Interestingly, among the downregulated transcripts, we identified multiple GO terms and gene sets associated with cell-cycle regulation, in agreement with the growth-inhibitory phenotype observed in TES-expressing cells (**Figure 2B,C, Figure S3C,D** and **Table S3**).

**Figure 2.**
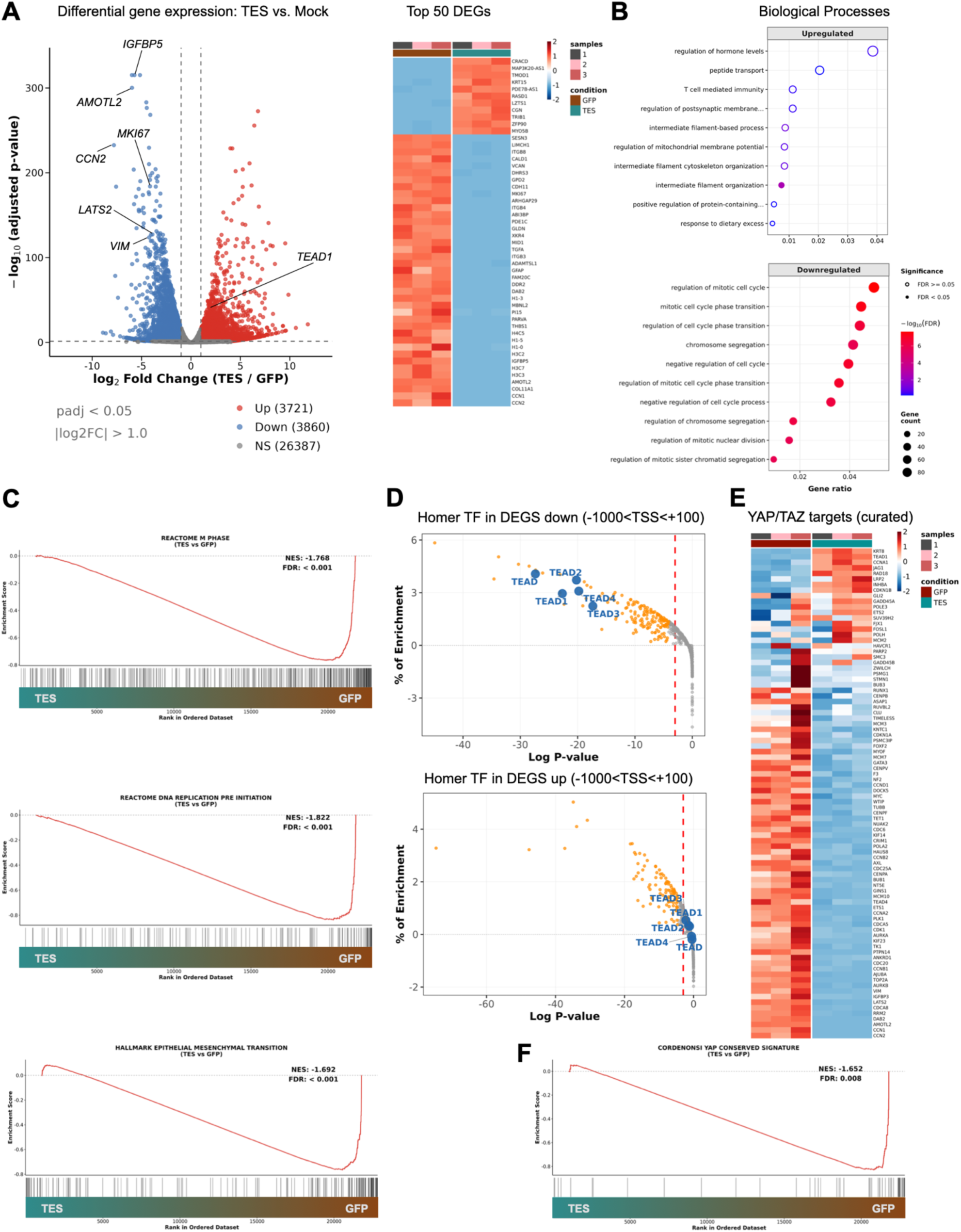
TES induces widespread transcriptional changes. (A) TES induces robust gene deregulation compared to mock cells two days after SNB19 infection, as assessed by RNA sequencing summarized as volcano plot (RNA-seq; padj < 0.05; |log_2_ Fold Change| > 1.0) and the heatmap of the top 50 differentially expressed genes (DEGs). (B) Gene Ontology (GO) analysis indicates that genes upregulated showed limited functional enrichment, whereas genes associated with cell cycle regulation are downregulated, as expected given the transcriptional repressive role of TES. FDR, false discovery rate. (C) Enrichment plots from gene set enrichment analysis (GSEA) for Reactome M phase, Reactome DNA replication pre initiation, and Hallmark epithelial mesenchymal transition (see also Table S3). NES, normalized enrichment score. (D) Homer analysis shows that transcription factor binding sites (TFBSs) belonging to the TEAD family were significantly enriched only among differentially expressed downregulated genes. TSS, transcription start site. (E) Heatmaps with 94 direct YAP/TAZ target genes, showing that the large majority of them are downregulated by TES. (F) Enrichment plot from GSEA for conserved YAP transcriptional signature described by Cordenonsi et al^43^.

Given that TES is expected to bind the same genomic targets as TEAD1, we next analyzed the enrichment of transcription factor binding sites (TFBSs) within promoter regions (−1000/+100 bp from the transcription start site) of differentially expressed genes using the HOMER algorithm^33^. Strikingly, TFBSs belonging to the TEAD family were significantly enriched among downregulated genes, whereas they were absent from the upregulated gene set (**Figure 2D**).

We then examined the expression of established TEAD1/YAP/TAZ target genes. To this end, we curated a list of 94 genes previously validated as direct YAP/TAZ targets through functional assays across multiple experimental systems^25,34–42^ (**Table S4**). The vast majority of these genes were downregulated following TES expression in SNB19 cells (**Figure 2E**). Consistently, GSEA performed using the conserved YAP transcriptional signature described by Cordenonsi et al.^43^revealed a strong negative enrichment in TES-treated cells (**Figure 2F**).

To further characterize TES activity at the chromatin level, we assessed its genome-wide occupancy using CUT&Tag approach^44^. We compared TES binding capability with that of wild-type TEAD1, both tagged (V5 for TES and HA for TEAD1) and overexpressed at comparable levels via LV transduction in SNB19 cells. Peak calling revealed partially distinct binding profiles for TES and TEAD1 (**Figure 3A, Figure S4A** and **Table S5**). However, in the majority of genomic regions where one factor was bound, the other was also detected, albeit often at lower signal intensity (**Figure 3B**). Peaks unique to TES, unique to TEAD1, or shared between the two displayed similar genomic distributions, with a large fraction localized to putative enhancer regions, as previously reported for TEAD factors^38^ (**Figure S4B**). Motif analysis confirmed the presence of TEAD-recognized DNA motifs in both TES and TEAD1 peak sets (**Figure 3C**). Together, these data indicate that TES largely preserves the wild-type TEAD1 DNA-binding specificity (**Figure 3B,C, Figure S4A-C**).

**Figure 3.**
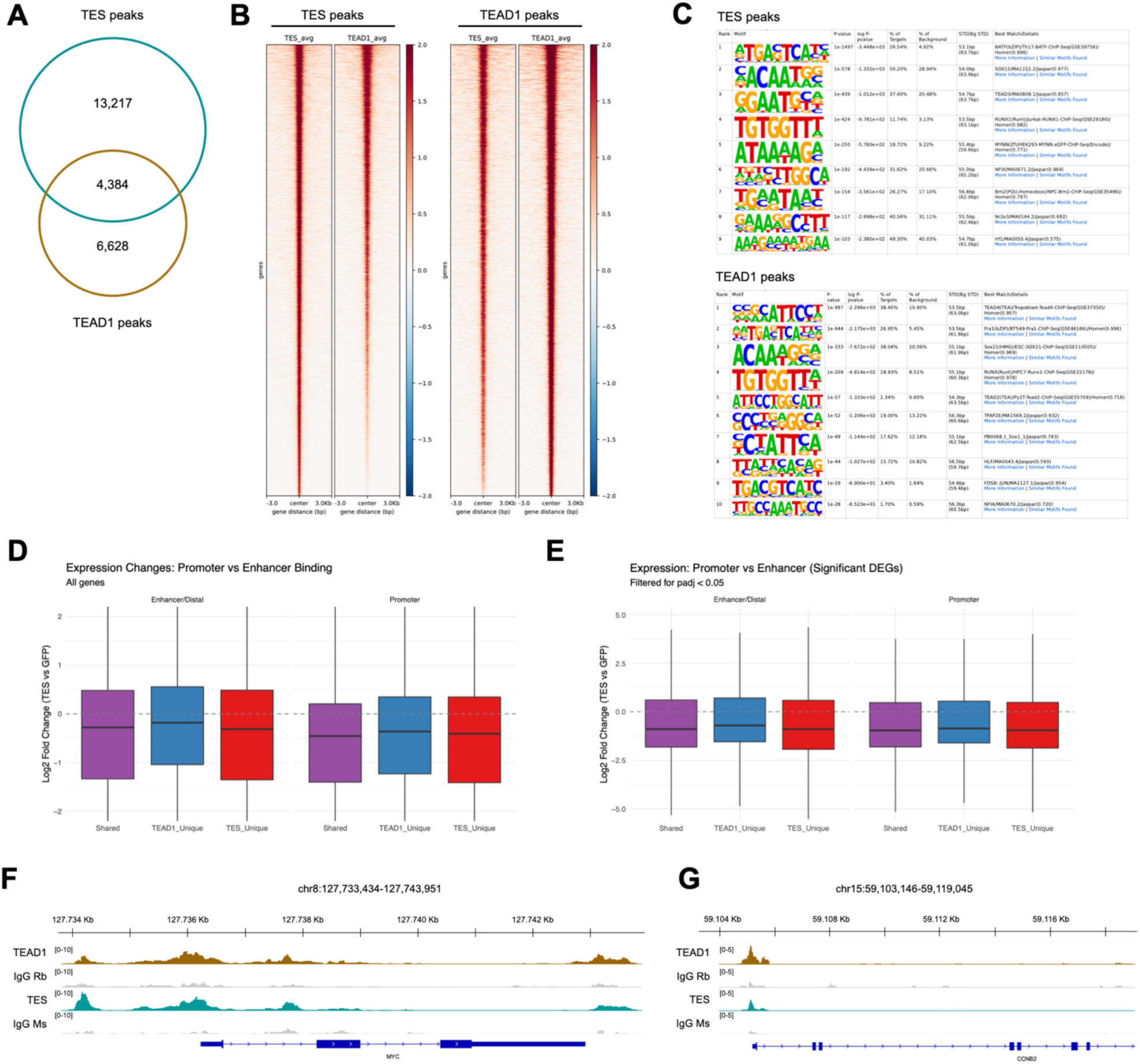
TES induces transcriptomic and epigenetic changes. (A) Venn Diagram showing the overlap between TES and TEAD1 peaks identified by CUT&Tag analysis in SNB19 cells. (B) Heatmaps showing relative enrichment for CUT&Tag (V5 tag for TES and HA tag for TEAD1 overexpression) in both control and TES condition. Enrichments are shown as color scale in peak bodies of ± 3 Kb. (C) Motif enrichment analysis confirming the presence of TEAD-recognized DNA motifs in both TES and TEAD1 peak sets. Motifs are ranked based on *P*-value, with corresponding log P-values, percentage of target sequences, and percentage of background frequency indicated. (D, E) Integration of TES genomic occupancy with transcriptional changes. CUT&Tag peaks for TES only, TEAD1 only or TES/TEAD1 shared were annotated as promoter-associated or enhancer-associated regions and intersected with all genes (D) or with differentially expressed genes (DEGs, padj < 0.05; E) identified by RNA-seq. The graphs show expression changes (log2 fold change) of genes associated with TES/TEAD1 binding sites at promoters or enhancers compared to mock condition (GFP). (F, G) IGV (Integrative Genomics Viewer) snapshots of *MYC* (F) and *CCNB2* (G) loci showing CUT&Tag tracks in TEAD1- (yellow tracks) and TES-overexpressing (blue tracks) cells. TES largely preserves TEAD1 DNA-binding specificity, as evidenced by the conserved occupancy profile at both *MYC* and *CCNB2* loci.

We next intersected TES genomic occupancy with transcriptional changes. Genes associated with TES/TEAD1 binding sites were generally downregulated in TES-expressing cells compared to GFP controls (**Figure 3D**), an effect that became more pronounced when considering only differentially expressed genes (**Figure 3E-G**).

Collectively, these data indicate that TES retains the DNA-binding activity of parental TEAD1 and induces transcriptional repression of its target genes.

### TES antitumor activity in mouse xenograft models

To investigate the antitumor activity of TES *in vivo*, we established heterotopic subcutaneous xenografts in non-obese diabetic–severe combined immunodeficient gamma (NSG) mice using patient-derived GBM CSCs previously transduced with lentiviruses expressing either GFP (mock) or TES (**Figure S5A**). Tumors developed in both conditions; however, xenografts derived from TES-expressing cells were consistently smaller than those generated from GFP-expressing controls, indicating that TES expression attenuated tumor growth (**Figure S5B**).

We next assessed TES activity in an orthotopic setting by intracranially transplanting GFP-or TES-expressing CSCs into NSG mice (**Figure 4A**). At both three and five weeks post xenotransplantation (WPX), tumors originating from TES-transduced cells were significantly smaller than those formed by control cells (**Figure S5C,D** and **Figure 4B,C**). Immunohistochemical analysis revealed that control tumors exhibited high proliferative activity, as indicated by abundant phosphorylated histone H3 (PH3)-positive cells, whereas TES-expressing tumors showed markedly reduced proliferation accompanied by extensive cell death, evidenced by strong cleaved caspase-3 (CC3) staining (**Figure S5E,F**).

**Figure 4.**
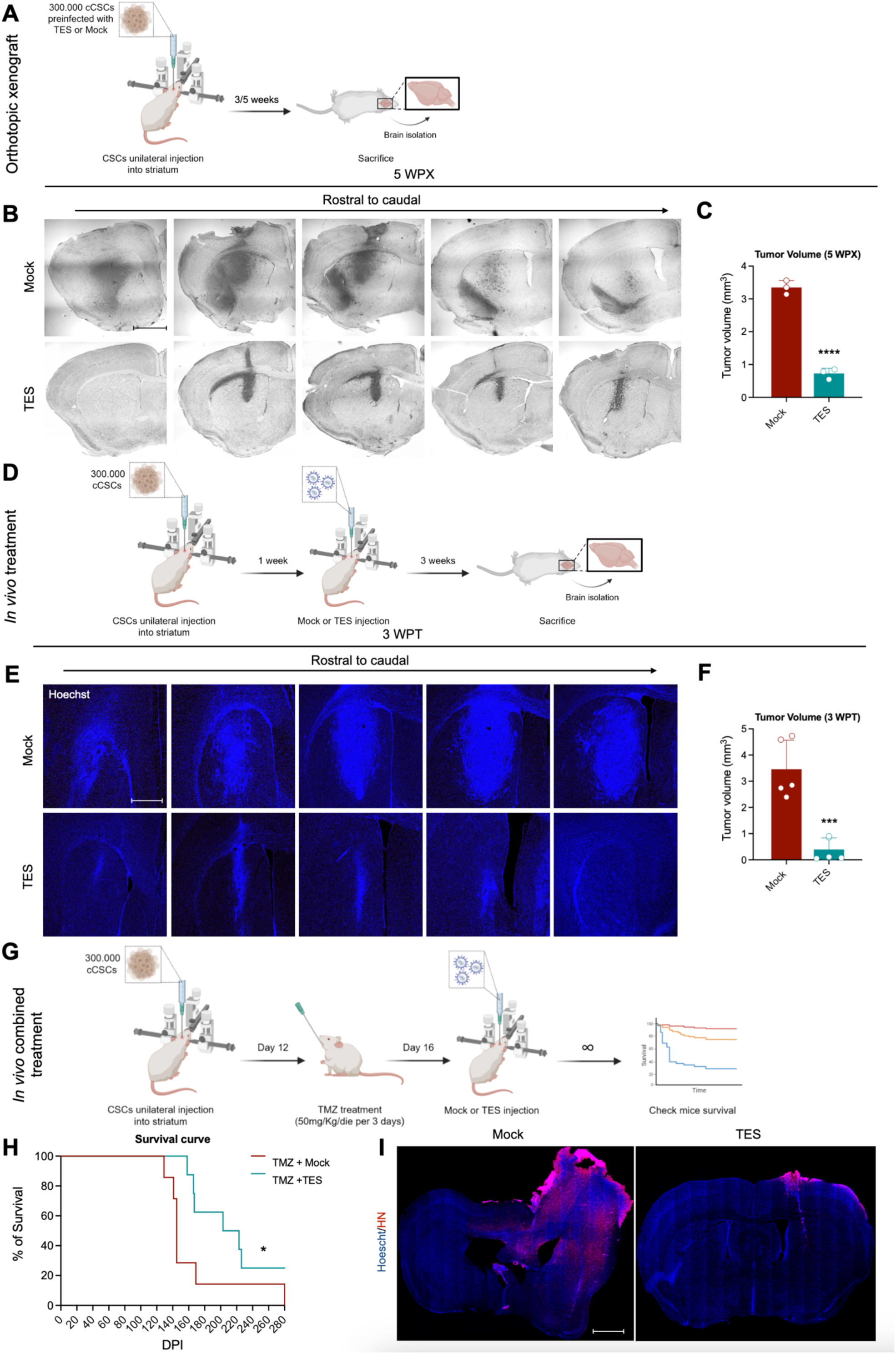
TES treatment restrains orthotopic tumor growth in immunodeficient mice. (A) Experimental scheme illustrating the generation of orthotopic xenografts by intracranial transplantation of Mock or TES pre-infected CSCs into the striatum of immunodefiicent NSG mice. (B) Nissl staining of representative sections of the orthotopic xenograft model, 5 weeks after pre-infected CSCs transplantation, showing strong ability of TES in inhibiting tumor growth compared to Mock. Scale bar: 400 μm. (C) Quantification of tumor volume: Mock vs TES \*\*\*\**P*<0.0001. Data are expressed as means ± SEM. Statistical analysis was performed using unpaired *t* test. *n* = 3 animals per group. WPX, weeks post xenotransplantation. (D) Experimental scheme illustrating the *in vivo* treatment of orthotopic xenografts in NSG mice. After 1 week from CSCs transplantation, local intratumoral injections of Mock or TES delivered by LV were performed directly into growing intracranial GBM xenografts. (E) Hoechst staining of representative sections of the described model in Fig. 4D, 3 weeks after local intracranial LVs injection, demonstrating the anti-tumor activity of TES treatment by strongly restraining tumor growth compared to Mock. Scale bar: 200 μm. (F) Quantification of tumor volume: Mock vs TES \*\*\**P*=0.0004. Data are expressed as means ± SEM. Statistical analysis was performed using unpaired *t* test. *n* = 5 animals per group. WPT, weeks post treatment. (G) Experimental scheme illustrating the *in vivo* combined treatment (TMZ + LV-TES) of orthotopic xenografts in NSG mice. After 12 days from CSCs transplantation, mice were treated with TMZ (50mg/Kg/die for 3 days) through oral gavage. After 1 day of wash-out from the treatment, animals were locally injected with LV expressing Mock or TES into the striatum. Mice were monitored for survival analysis and euthanized upon observation of signs of reduced general health. (H) Survival curve analysis reports an increase of around 2 months of the overall survival of mice treated with TMZ+LV-TES compared to TMZ+Mock: Mock vs TES \**P*=0.0207. Data are expressed as percentage of survival. Statistical analysis was performed using Gehan-Breslow-Wilcoxon test. *n* = 7 animals for TMZ+Mock group, *n* = 8 animals for TMZ+TES group. DPI, days post injection. (I) Immunohistochemistry on representative coronal sections for Human Nuclei (HN, marker for human tumor cells transplanted) counterstained with Hoechst, of Mock and TES mice at the endpoint (280 days from cells transplantation). HN staining shows that combined TMZ and TES treatment significantly restrains tumor mass growth in mice. Scale bar: 500 μm.

To investigate the therapeutic potential of TES in an established tumor context, we performed local intratumoral lentiviral delivery directly into growing intracranial GBM xenografts. CSCs were allowed to engraft and form detectable tumors for seven days prior to intracranial injection of either GFP- or TES-expressing LV (**Figure 4D**). Analysis three weeks post treatment (WPT) revealed a pronounced reduction in tumor growth in mice treated with TES-LV compared to controls (**Figure 4E,F**). Given that temozolomide (TMZ) represents the current standard-of-care chemotherapy for GBM, we next evaluated the therapeutic efficacy of TES in combination with TMZ. Tumor-bearing mice were treated with TMZ for three consecutive days prior to lentiviral injection (**Figure 4G**). Combined TMZ and TES treatment significantly improved animal survival compared to TMZ alone, extending median survival by approximately two months and resulting in ∼20% long-term survival rate (**Figure 4H**). At the experimental endpoint (280 days), mice receiving combined TES and TMZ treatment, while not tumor-free, displayed markedly reduced tumor burdens (**Figure 4I**).

Together, these data demonstrate that TES exerts robust antitumor activity *in vivo*, significantly reducing tumor growth and enhancing therapeutic efficacy in GBM xenograft models.

### TES impairs cell migration

Since TEAD1 has been reported as regulator of mobility and mesenchymal transformation in GBM^24^, we intended investigate whether TES expression could regulate this aspect. GSEA analysis of transcriptome of TES-treated cells revealed a downregulation of migration-associated gene sets (**Figure S6A**). To validate experimentally the activity of TES against cell migration, we firstly assessed whether TES treatment impairs wound healing of SNB19 cells (**Figure 5A**). We found that TES treatment impaired the capability of the cells to fill the scratch within the 48 hrs that not infected cells needed (**Figure 5B**). Secondly, we adopted the transwell invasion assay (**Figure S6B**). Even with this protocol, we confirmed that TES is inhibiting the migration of the SNB19 cells (**Figure S6C**). Lastly, we performed the spheroid dispersion assay using tumorspheres deriving from patient-derived CSCs (mesenchymal subtype) previously infected with LVs. Comparing the area of migration at the plating (t 0) and after three days (t 72 hrs), we found that TES is affecting the sprouting from the spheres (**Figure S6D**). Importantly, (i) TES effect is similar to the effect of YAP/TAZ inhibitor verteporfin in both wound healing and transwell assays^45^ (**Figure 5B** and **Figure S6C**), (ii) it was absent using the mutated TES^32^ (**Figure 5B** and **Figure S6C,D**), and (iii) the experiments have been performed when the proliferation differences are not yet evident (**Figure S1**).

**Figure 5.**
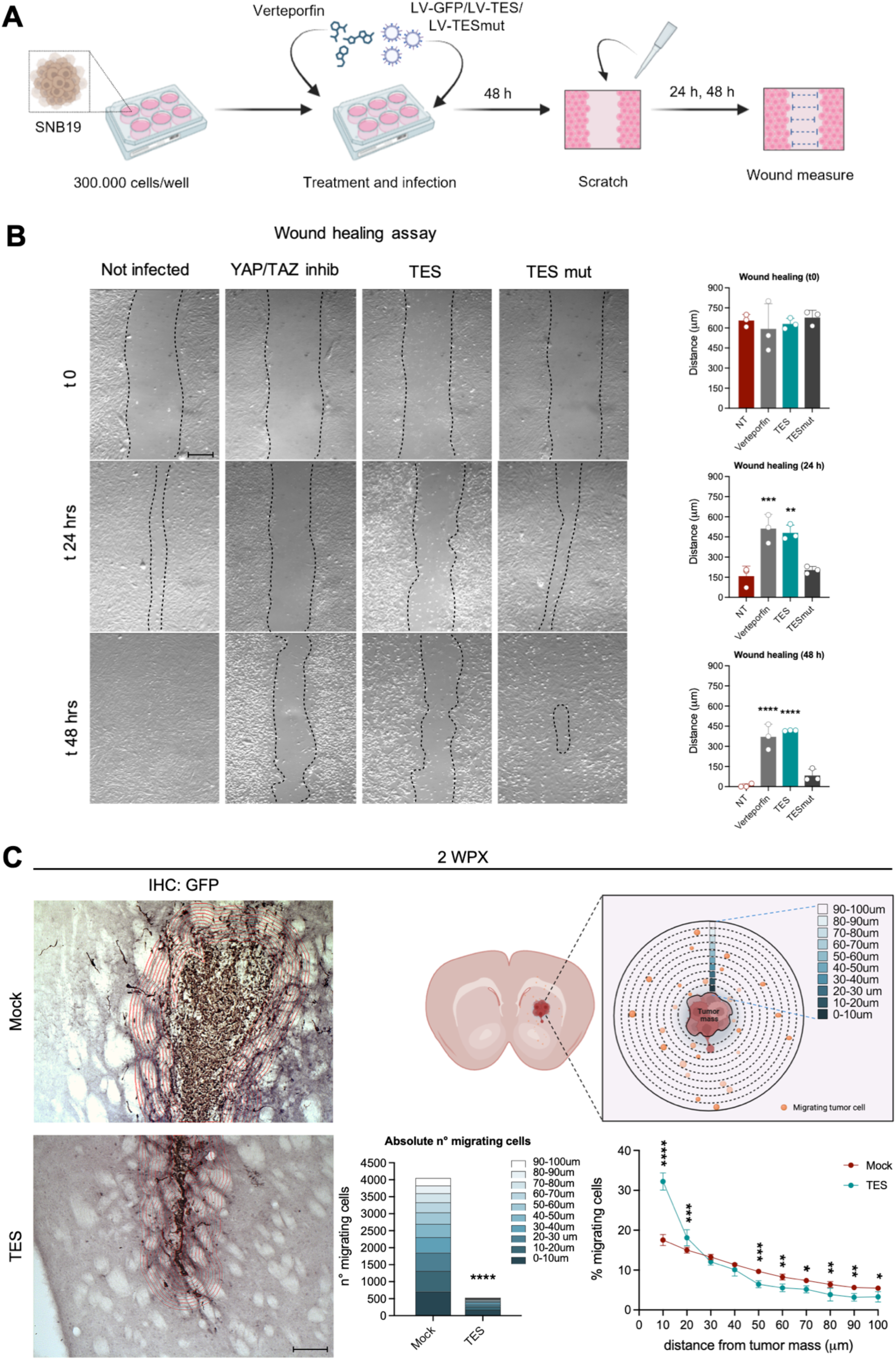
**TES reduces tumor cell migration both *in vitro* and *in vivo*** (A) Experimental scheme of the treatment protocol used in SNB19 cell line to perform the wound healing assay. (B) Representative bright-field images and wound closure measures of SNB19 not infected (NT), treated with Verteporfin (YAP/TAZ inhibitor, 0.2 μM), infected with LV-TES and with LV-TESmut, at t0, 24 and 48 hours after scratch, revealing TES capability of impairing cells ability to fill the scratch. Wound closure distance t0: NT vs Verteporfin *P*=0.7988; NT vs TES *P*=0.9821; NT vs TESmut *P*=0.9871. Wound closure distance t24: NT vs Verteporfin \*\*\**P*=0.0008; NT vs TES \*\**P*=0.0015; NT vs TESmut *P*=0.7791. Wound closure distance t48: NT vs Verteporfin \*\*\*\**P*<0.0001; NT vs TES \*\*\*\**P*<0.0001; NT vs TESmut *P*=0.2647. Data are expressed as means ± SEM. Statistical analysis was performed using ordinary one-way ANOVA followed by Dunnett’s multiple comparisons test. *n* = 3. Scale bar: 300 μm. (C) TES treatment in orthotopic mouse xenografts (represented in Fig. 4A) strongly limits tumor cell invasion into peritumoral tissue at 2WPX. On the left, immunohistochemistry on representative coronal brain sections for GFP (tag for tumor cells) in animals injected with Mock or TES pre-infected GFP-labelled CSCs at 2 WPX. Tumor cell migration from the primary tumor mass was assessed by manually counting migrating cells within 10-μm concentric regions of interest (ROIs) delineated around the tumor core at increasing distances (0–100 μm). 10-μm concentric ROIs marked by red dash line in representative immunohistochemical images are shown. On the right, quantification of tumor cell migration reported as absolute number of migrating cells for each distances (Mock vs TES \*\*\*\**P*<0.0001) and as percentage of migrating cells for each distances normalized to the total number of migrating cells (Mock vs TES 0-10 μm \*\*\*\**P*<0.0001; Mock vs TES 10-20 μm \*\*\**P*=0.0004; Mock vs TES 20-30 μm *P*=0.6342; Mock vs TES 30-40 μm *P*=0.5606; Mock vs TES 40-50 μm \*\*\**P*=0.0002; Mock vs TES 50-60 μm \*\**P*=0.0025; Mock vs TES 60-70 μm \**P*=0.0243; Mock vs TES 70-80 μm \*\**P*=0.0052; Mock vs TES 80-90 μm \*\**P*=0.0078; Mock vs TES 90-100 μm \**P*=0.0280). Data are expressed as means ± SEM. Statistical analysis was performed using two-way ANOVA followed by Sidak’s multiple comparisons test. *n* = 5 animals per group. Scale bar: 100 μm.

To confirm whether the anti-migratory properties observed in cell culture are recapitulated *in vivo*, we performed orthotopic xenotransplantation as performed above (**Figure 4A**) using CSCs preinfected with GFP-LV with or without TES-LV coinfection. Histochemical analysis at 2 WPX for GFP^+^ cell distribution on brain slices containing the tumor evidenced differences in infiltrative tumor spread (**Figure 5C**). While control cells had infiltrated in the striatum, TES-treated cells were found very close to the tumor edge (**Figure 5C**).

These findings suggest that beyond the effect on cell proliferation TES treatment has additional outcome on GBM cells reducing their infiltrative capacity.

### TES is safe for normal brain cells

YAP/TAZ signaling plays essential roles during neural development, whereas it is largely inactive in mature brain cells^46^. Therefore, repression of TEAD1-dependent transcription by TES in the adult brain is expected to be intrinsically well tolerated. To directly assess the effects of TES on healthy neural cells, we first evaluated its impact on primary mouse cortical neuron cultures. Neurons were transduced with LVs expressing either TES or GFP, and cell survival was analyzed three weeks later (**Figure 6A**). In both conditions, neurons displayed comparable morphology, as assessed by MAP2 immunostaining, and similarly low numbers of propidium iodide–positive (PI⁺) cells, indicating no increase in cell death upon TES expression (**Figure 6B**).

**Figure 6.**
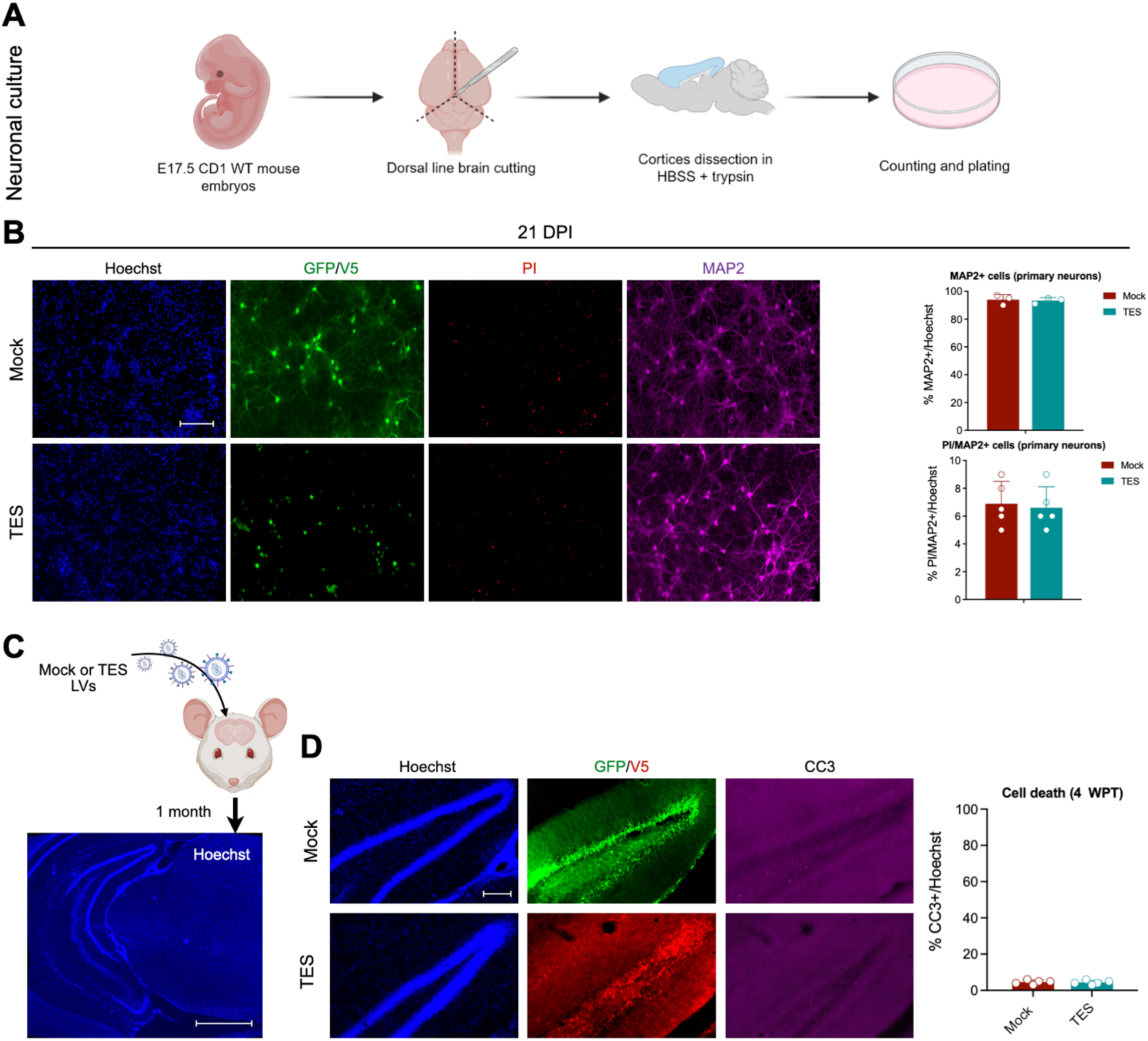
**TES expression does not affect healthy brain cells both *in vitro* and *in vivo*** (A) Experimental scheme of the dissection protocol performed to obtain primary mouse neuronal cultures from mouse embryonic cortices of E17.5 (embryonic day 17.5) CD1 WT mice. (B) Immunocytochemistry for GFP (mock tag), V5 (TES tag), propidium iodide (PI, cell death) and MAP2 (neuronal marker) counterstained with Hoechst of primary murine cortical neurons 21 days after the infection with either mock or TES-expressing lentiviruses, indicating no specific neuronal toxicity induced by TES. (*Upper)*, quantification as percentage of MAP2+ cells on the total number of Hoechst+ nuclei in Mock or TES infected neurons at 21DPI, demonstrating maintenance of primary neurons population after the treatment: Mock vs TES *P*=0.7953. *n* = 3. (*Lower)*, quantification as percentage of PI+ cells on the total number of MAP2+/Hoechst+ nuclei in Mock or TES infected neurons at 21DPI, reporting no increase in cell death after TES treatment: Mock vs TES *P*=0.7684. *n* = 5. Data are expressed as means ± SEM. Statistical analysis was performed using unpaired *t* test. Scale bar: 200 μm. (C) Experimental scheme illustrating the intracranial local injection of TES- or GFP-expressing lentiviruses into the hippocampus of adult wild-type C57BL/6 mice. (D) Immunohistochemistry on representative coronal brain sections for GFP (Mock tag), V5 (TES tag) and CC3 (cell death marker) counterstained with Hoechst of animals injected with Mock or TES at 4WPT. Quantification of CC3+ cells within infected hippocampi shows no difference between TES and mock treated animals, indicating that TES is not toxic for murine healthy brain cells: Mock vs TES *P*=0.7885. Data are expressed as means ± SEM. Statistical analysis was performed using unpaired *t* test. *n* = 5 animals per group. Scale bar: 100 μm.

We next examined TES safety *in vivo* by stereotactically injecting TES- or GFP-expressing lentiviruses into the hippocampus of adult wild-type C57BL/6 mice (**Figure 6C**). Injected animals were monitored for one month following surgery, during which no overt behavioral abnormalities were observed. Consistently, histological analysis revealed comparable levels of cell death (CC3^+^ cells) in brains injected with either TES or GFP control vectors (**Figure 6C**).

Together, these findings indicate that TES expression does not adversely affect the survival or integrity of healthy neurons *in vitro* or *in vivo*, supporting the safety of TES for localized delivery in the adult brain.

## DISCUSSION

Glioblastoma remains a largely incurable disease, characterized by extensive intratumoral heterogeneity, diffuse infiltration into the surrounding brain parenchyma, and inevitable recurrence despite aggressive multimodal therapy. High YAP/TAZ activity correlates with poor prognosis and increased stemness, and YAP/TAZ effectors promote transcriptional programs supporting cell cycle progression and migration^43,37^. Given the complexity of this axis, direct inhibition of YAP/TAZ/TEAD interactions has seen limited clinical translation. Small molecules targeting TEAD palmitoylation or YAP–TEAD interfaces have shown promise *in vitro* but face challenges in potency, specificity, and delivery across the blood–brain barrier^47,48^. These limitations have spurred interest in alternative strategies that rewire, rather than simply block, aberrant transcriptional networks.

Transcription factor–based therapies (e.g. delivering reprogramming factors or dominant-negative TFs) have been proposed to shift tumor cells out of proliferative or stem-like states. However, direct overexpression of TFs runs the risk of off-target effects and lacks mechanisms for durable suppression of pathological gene networks. In contrast, engineered transcriptional repressors and epigenetic modulator fusions (epieditors) offer a programmable avenue to modulate entire gene programs by imposing stable chromatin changes at endogenous loci^18–21^.

In this study, we introduce a novel epigenetic silencer factor (ESF) based on the TEAD1 transcription factor and demonstrate its ability to suppress YAP/TAZ-dependent transcriptional programs, impair tumor growth, and enhance therapeutic efficacy in GBM models, while sparing healthy brain tissue. Our work builds upon the concept that targeting single oncogenes or signaling nodes is insufficient to counteract the plasticity and adaptability of GBM cells, particularly cancer CSCs^15,49^. Instead, TES was designed to repress the entire transcriptional output downstream of YAP/TAZ. By retaining the DNA-binding specificity of TEAD1 while converting it into a stable transcriptional repressor, TES effectively hijacks an oncogenic transcriptional platform and rewires it toward durable gene silencing.

Transcriptomic analyses revealed that TES induces widespread transcriptional repression, with a strong bias toward downregulation of coherent gene sets associated with cell-cycle progression and proliferation. This effect is consistent with the observed reduction in glioma cell growth and induction of cell death across established glioma cell lines and patient-derived CSCs. These results are consistent with previous observations that YAP/TAZ drives a pro-growth program in GBM and other cancers^43,38^. In addition to its anti-proliferative activity, TES exerted a clear anti-migratory effect, in line with extensive evidence implicating TEAD factors as key drivers of GBM cell migration and invasiveness^24^.

Both enrichment analyses, motif discovery and genome-wide occupancy profiling further showed that TES largely mirrors the chromatin-binding landscape of wild-type TEAD1 and selectively represses YAP/TAZ target genes. This observation is consistent with previous genome-wide studies showing that TEAD transcription factors recognize a highly conserved DNA motif and act as the primary chromatin-anchoring platform for YAP/TAZ-driven transcriptional programs^38,50^. In multiple cellular contexts, TEAD factors have been shown to occupy regulatory regions independently of YAP/TAZ, which are instead required to convert TEAD-bound loci into transcriptionally active states^51,38^. Therefore, the selective repression of TEAD motif–containing genes by TES supports a model in which TES exploits this intrinsic DNA-binding specificity to convert an oncogenic transcriptional hub into a repressive platform, rather than inducing nonspecific transcriptional toxicity.

Functionally, TES exhibited strong antitumor activity *in vivo*. In both heterotopic and orthotopic xenograft models, TES expression significantly reduced tumor growth and proliferation while promoting tumor cell death. Importantly, TES was effective not only when expressed prior to transplantation, but also when delivered locally into established intracranial tumors, supporting its potential therapeutic relevance. Moreover, combination treatment with temozolomide resulted in a substantial survival benefit compared to chemotherapy alone. These results suggest that TES may sensitize GBM cells to standard therapy or counteract resistance mechanisms that limit TMZ efficacy.

A major concern for any transcriptional or epigenetic therapy targeting the brain is potential toxicity to normal neural cells. YAP/TAZ signaling is largely inactive in the adult brain, being primarily engaged during development and in regenerative contexts. Consistently, TES expression did not adversely affect primary neurons *in vitro* nor induce detectable toxicity or behavioral alterations when delivered to the adult mouse brain *in vivo*. While more extensive safety studies will be required, these findings define a favorable therapeutic window and support the feasibility of localized TES delivery in the CNS.

GBM progression and recurrence are sustained by the coexistence of multiple transcriptional states, including proliferative, stem-like, invasive, and therapy-resistant phenotypes. Targeting a single pathway is therefore unlikely to achieve durable tumor control, whereas coordinated repression of parallel or convergent transcriptional networks may be required to overcome tumor plasticity.

In this context, TES-based repression of YAP/TAZ-TEAD–dependent transcription could be synergistically combined with other therapies that target orthogonal vulnerabilities. For example, oncolytic viruses or suicide gene therapies^52^ could benefit from TES-mediated suppression of survival and stress-response programs, potentially enhancing tumor cell susceptibility to cytotoxic insults. Similarly, TES could be combined with immunomodulatory gene therapies, as YAP/TAZ signaling has been implicated in shaping an immunosuppressive tumor microenvironment and promoting immune evasion in multiple cancers^53,54^. Repressing YAP/TAZ-dependent programs may therefore improve immune infiltration or sensitize tumors to immune-based interventions.

Notably, TES is conceptually well suited for combination with other ESFs designed to target distinct transcriptional hubs. In particular, our previously developed SOX2 Epigenetic Silencer (SES)^17^. While SES primarily disrupts lineage-specific and stem-cell–associated gene networks, TES suppresses YAP/TAZ-driven programs linked to proliferation, invasion, and therapy resistance. Because both TES and SES act using endogenous DNA-binding platforms through epigenetic repression at physiologically relevant regulatory elements, their combined use could impose multi-layered and durable transcriptional constraints that are difficult for tumor cells to bypass through genetic or epigenetic adaptation. This strategy may be particularly effective in preventing tumor recurrence, which is often driven by residual CSCs that dynamically switch between transcriptional states in response to therapy.

Together, our results position TES as a proof-of-concept for a broader class of epigenetic silencer–based therapies aimed at rewiring oncogenic transcriptional networks rather than inhibiting single molecular targets. This strategy may be particularly well suited for highly plastic and heterogeneous tumors such as GBM, where durable suppression of stemness, proliferation, and invasiveness is required to prevent recurrence. Future work will focus on optimizing TES delivery, and exploring combination strategies with radiotherapy, immunotherapy, or additional pathway-targeting ESFs. Ultimately, expanding the ESF toolkit may enable customizable, pathway-level interventions tailored to the molecular dependencies of individual tumors.

## MATERIALS AND METHODS

### Constructs

***KRAB-hTEAD1_1-166_-DNMT3A/3L (TES).*** The construct was designed using the engineering strategy previously described by our laboratory (Benedetti et al., 2022). Notably, the initial part of human *TEAD1* gene, coding for amino acids 1 to 166 (thus excluding the TEAD1 activator domain) was fused with the KRAB repressor domain (from the gene *ZNF10* encoding for a zinc finger protein; amino acids 1 to 97) at its N-terminus, whereas the functional domains of DNMT3A (amino acids 388 to 689) and DNMT3L (amino acids 206 to 421) were fused at the C-terminus of theTEAD1 portion. The V5 tag was fused at the end of the new chimeric transgene. The transgene was inserted in a lentiviral construct, with Ef1αL as promoter.

***TESmut (R85K).*** TES construct was mutated in the residue 85 of the initial part of human *TEAD1* gene (arginine in position 85 to lysine). This R85K mutation might alter the binding affinity or specificity of TEAD1 to its DNA targets.

GFP expressing lentiviruses were used for all the following experiments as control condition (Mock), exploiting the same Ef1αL promoter for protein expression.

### Lentivirus production

To generate lentiviruses (LV), lentiviral replication-incompetent particles coated with the vesicular stomatitis virus glycoprotein (VSVg) were packaged in HEK293T cells. Cells were transfected with 32 μg of vector and packaging constructs, according to conventional CaCl2 transfection protocol. After 30 hours, medium was collected, filtered through 0.44μm cellulose acetate and centrifuged at 20,000 rpm for 2 hours at 20°C in order to concentrate the virus. Each viral batch was titrated with a dedicated kit (Applied Biological Materials, LV900). The titer obtained is in IFU/ml (infections units per ml) and, according to the manufacturer instructions, the amount of genomic RNA in a sample can be converted to its viral titer by calibration with the standards of lentiviral preparations provided. The viral batch was used when titer was 10^9 IFU/ml. For all the *in vitro* experiments a MOI of 33 was used.

### Cell cultures

U87 and SNB19 (human glioma cell lines) were cultured in plastic-adherence condition in DMEM medium (Dulbecco’s Modified Eagle’s Medium - high glucose, Sigma-Aldrich) containing 10% fetal bovine serum (FBS, Sigma-Aldrich), 1% penicillin/streptomycin (Pen/Strept, Sigma-Aldrich), 2mM Glutamine (Sigma-Aldrich), 1% non-essential amino acids (MEM NEAA, ThermoFisher Scientific), 1% sodium pyruvate solution (Sigma-Aldrich). All the cell lines were passaged twice a week using a trypsin-EDTA solution (Sigma-Aldrich). Cancer stem cells (CSCs, gift of Dott.ssa Rossella Galli) derived from patients with classical (L0627) and mesenchymal (L1312) glioblastoma tumors were maintained in spheres in suspension cultures in DMEM/F12 medium (Sigma-Aldrich) supplemented with Hormon Mix (DMEM/F12, 0.6% Glucose (Sigma-Aldrich) (30% in phosphate buffer (PBS, Euroclone), 250 μg/ml insulin (Sigma-Aldrich), 97 μg/ml putrescine powder (Sigma-Aldrich), 1 mg/mL apotransferrin powder (Sigma Aldrich), 0,3 μM sodium selenite, 0,2 μM progesterone), 1% Pen/Strept, 2mM Glutamine, 0.66% Glucose (30% in PBS), and 4mg/ml heparin (Sigma-Aldrich); basic fibroblast growth factor (bFGF, 20 ng/ml, ThermoFisher Scientific) and epidermal growth factor (EGF, 20 ng/ml, ThermoFisher Scientific) were freshly added to culture medium every 2 days. Sphere cultures were passaged once a week by mechanical dissociation of the spheres to single cell suspension. All the cell cultures were kept in humidified atmosphere of 5% CO2 at 37°C under atmospheric oxygen conditions. All the cell lines used in this work are listed in Table S6.

### Cell growth analysis

Two different protocols for the cell growth analysis were exploited according to the specific culturing conditions.

For adhesion cell lines: 1,2×10^5 cells/well (SNB19 and U87) were seeded in adherent condition in a 12-multiwell plate; the following day (day 0), cultures were infected with lentiviral vectors (4 µl/well of LV 1×10^9 IFU/mL). At day 3 cells were detached, and live cells were stained with Trypan Blue Solution (0.4%, ThermoFisher Scientific) and counted using CountessTM II Automated Cell Counter (ThermoFisher Scientic); after this passage, 1,2×10^5 cells were reseeded until the cell number obtained was no longer sufficient to allow plating at this density. This procedure was repeated every 3 days, while the overall experiment was repeated 3 times for each time point. Brightfield representative images were taken at each time-point.

For suspension cell lines: 3×10^5 CSCs/well (both L0627 and L1312) were infected with lentiviral vectors (10 µl/well of LV 1×10^9 IFU/mL) and seeded in single cell suspension condition in a 6-multiwell plate at day 0. At day 4, day 7 and day 11 CSCs spheres were mechanically dissociated to single cell suspension, and live cells were stained with Trypan Blue Solution and counted as previously described. Live cell number and the % of dead cell were reported on graphs for each time point; the experiment was repeated 3 times for each time-point. Bright-field representative images were taken at each time-point. Replicates represent the number of cells counted in independent plating.

### Western blot analysis

5×10^5 SNB19 cells were plated in each well of a 6-multiwell plate. The following day (day 0) the cells were infected with lentiviruses (17 μl/well of LV 1×10^9 IFU/mL). At day 2, cells were homogenized in RIPA buffer (50 mM Tris pH 7.5, 150 mM NaCl, 1 mM EDTA, 0,1 % SDS, 1% Triton X-100, Roche Complete EDTA-free Protease Inhibitor Cocktail, Roche PhosSTOP EASYpack) and proteins were extracted. Protein lysates were quantified using the Pierce BCA Protein Assay Kit (Thermo Fisher Scientific). 10 μg of proteins were loaded in Laemmli buffer 4X on a polyacrylamide gel and then separated in the running gel at 150-mV electric field. To detect our proteins, a 4%-12% polyacrylamide gels were used. The proteins were then transferred from the gel to a nitrocellulose membrane using the Trans-Blot Turbo transfer system (BioRad). At the end of the transfer, the membranes were stained with Red Ponceau (0.1% of 5% acetic acid solution and 0.01% Red Ponceau in ultrapure H2O) to visualize the protein bands, and then incubated for one hour at room temperaure in a 5% w/v solution of bovin serum albumine (BSA, Sigma-Aldrich) in Tris-Buffered Saline 0.1% Tween (TBS-T, Merck) according to antibody datasheet, in order to saturate the nonspecific antibody binding sites. Afterwards, the membrane was incubated (overnight at 4°C) in the blocking solution supplemented with the primary antibody, selected according to the protein that required detection. The primary antibodies used were: anti-V5 (mouse, 1:1000, ThermoFisher Scientific, R96025) and anti-calnexin (rabbit, 1:2000, Sigma-Aldrich, C4731). The following day, the membranes were incubated with the corresponding species of secondary horseradish peroxidase (HRP)-conjugated antibody (diluted 1:5000 in 5% w/v BSA in TBS-T 1X) for 1 hour at room temperature. Immunodetection was performed using the Bio-Rad ChemiDoc Touch Image System instrument by exploiting the chemiluminescence reaction of SuperSignal West Pico PLUS Chemiluminescent Substrate ECL (Thermo Fisher Scientific, 34580) according to the manufacturer’s instructions. Band densitometry relative to control was calculated using ImageJ/Fiji software (National Institutes of Health, USA), normalized on housekeeping proteins. Antibodies used in this work are listed in Table S7.

### RNA isolation and real time qPCR

For RNA isolation, 5×10^5 SNB19 were plated in each well of a 6-multiwell plate, and the following day (day 0) the cells were infected with lentiviruses (17 μl/well of LV 1×10^9 IFU/mL). Cells were harvested at 2 days post-infection to extract the RNA as following. RNA was extracted using the TRIzol reagent isolation system (Sigma-Aldrich) according to the manufacturer’s instructions. For quantitative reverse transcription polymerase chain reaction (RT-qPCR), 1 μg of RNA was reverse-transcribed using the ImProm-II Reverse Transcription System (Promega). Obtained cDNA was diluted 1:10 and was amplified in a 20 μl reaction mixture containing 2 μl of diluted cDNA, 1× Titan Hot Taq EvaGreen qPCR Mix (Bioatlas) and 0.6 μM of each primer. RT-qPCR was performed in triplicate with custom designed oligos using the CFX96 Real-Time PCR Detection System (Bio-Rad, USA). *GAPDH* transcripts were used as reference controls. Analysis of relative expression was performed using the ΔΔCt method and the CFX Manager software (Bio-Rad, USA). Primers used in this work are listed in Table S8.

### *In vitro* migration assays

Migratory behavior was assessed using wound healing, transwell migration and sphere diffusion assays. For wound healing and transwell migration assays, 3×10^5 SNB19/well were seeded in adherent condition in a 6-multiwell plate; the following day (day 0), cells were infected with lentiviral vectors (20 µl/well of LV 1×10^9 IFU/mL) or treated with 0,2 uM Verteporfin (YAP/TAZ inhibitor), and starting from day 1 cells were maintained in serum-free medium. Regarding the wound healing assay, two days post-infection a scratch (the wound) was created in the cell monolayer using a sterile pipette tip, and the capability of cells of migrating to heal the wound was evaluated after 24 and 48 hours. Bright-field images of the wound area were taken immediately (0 h) and at indicated time points, and the wound closure was quantified by measuring the remaining wound area using ImageJ/Fiji software. For transwell migration assays, two days post-infection, cells were detached and counted.

A total of 3×10^4 live cells were seeded into the upper chamber of transwell inserts (6.5 mm Transwell® with 8.0 µm Pore Polycarbonate Membrane Insert, Sterile, Merck, CLS3422) in serum-free medium and allowed to migrate through the 8µm pore-size membrane toward chemoattractant medium (medium supplemented with 1% of FBS) placed in the bottom chamber. After 24h migration was stopped by fixing the migrating cells with cold Methanol, followed by staining with Hoechst (1:1000 in PBS). Cell migration was measured by quantitating the number of migrating cells (adherent to the bottom of the membrane) per fields. Images were acquired with an epifluorescence microscope Nikon DS-Qi2 and analyzed with ImageJ/Fiji software. Regarding the sphere diffusion assay, mCSCs L1312 spheres were grown in low adherent condition for 1 week, and then 5 spheres/well were plated in a 24-multiwell and transduced with lentiviral vectors (10 µl/well of LV 1×10?^9 IFU/mL) in serum-free medium without EGF and bFGF growth factors. The following day, one sphere per condition was placed in poly-L-lysine (PLL, 10 μg/mL) + laminin substrate (5 μg/mL) in serum-free conditions without growth factors. Cell migration was assessed at 1 h (t0) and 72 h by measuring the area of confluent migration. To study the migration dynamics, bright-field images were taken at the indicated time points, and the diffusion area was quantified with ImageJ/Fiji software. Replicates represent the number of cells counted in independent platings. All the experiments were repeated at least 3 times.

### RNA-sequencing

RNA sequencing (RNA-seq) libraries were generated from 1 µg of total RNA isolated from SNB19 cells. RNA quantity and quality were assessed using a TapeStation instrument (Agilent Technologies), and only samples with an RNA integrity number (RIN) > 8 were used for library preparation to minimize 3′-end bias. RNA was processed according to the TruSeq Stranded mRNA Library Prep Kit protocol (Illumina), following the manufacturer’s instructions. Libraries were quantified, pooled, and sequenced on an Illumina HiSeq 3000 platform to generate 76-bp stranded paired-end reads using Illumina TruSeq technology. Image processing and base calling were performed using Illumina Real-Time Analysis software.

### CUT&Tag sequencing

CUT&Tag was performed according to the well-established CUTANA™ CUT&Tag Protocol (EpyCypher). Briefly, after obtaining a single-cell suspension for each experimental condition, cells were counted, and nuclei were extracted. 100.000 nuclei were used for each experimental replicate (three biological replicates for each condition). Afterward, nuclei were bound to activated concanavalin A-Conjugated Paramagnetic Beads (EpyCypher) and then incubated overnight at 4°C in lateral rotator with a primary antibody (V5 mouse, Thermo Fisher Scientific R96025; HA-tag rabbit, Cell Signaling Technology, 3724S) or control antibody (mouse, immunoglobulin G, Diagenode, C15400001; rabbit, immunoglobulin G, Diagenode, C15410206). The next day, nuclei suspensions were incubated with secondary antibody (Goat Anti-Mouse IgG Antibody (H+L), Unconjugated, Vector Laboratories, I-9200-1.5; Goat Anti-Rabbit IgG Antibody (H+L), Unconjugated, Vector Laboratories AI-1000-1.5), washed and DNA was fragmented. Chromatin tagmentation was performed using the pA-Tn5 Transposase (Diagenode, C01070001). DNA was then released from the nuclei, and sequencing libraries were amplified using a single-index barcode and then cleaned up. Finally, each individual library was paired-end sequenced on an Illumina NovaSeq platform.

### RNA-seq analysis

***Quality Control and Preprocessing.*** Raw sequencing reads were assessed for quality using FastQC. Adapter sequences and low-quality bases were removed using Trim Galore with the following parameters: Phred33 quality encoding (–phred33), quality threshold of Phred 20, minimum read length of 20 bp, adapter detection stringency of 3 bp (–stringency 3), and automatic Illumina adapter detection for paired-end reads (–paired). FastQC was also run on the trimmed reads to confirm successful adapter removal and quality improvement (–fastqc flag in Trim Galore). RNA-seq data generated in this study are accessible through the GEO database with the GEO series accession number GSE319888.

***Read Alignment.*** Trimmed reads were aligned to the human reference genome (GRCh38) using STAR aligner in two-pass mode (–twopassMode Basic). Key alignment parameters included: splice junction filtering (–outFilterType BySJout), maximum 20 multi-mapping locations (–outFilterMultimapNmax 20), maximum mismatch rate of 4% per mapped read length (–outFilterMismatchNoverReadLmax 0.04) with no hard cap on mismatch count (–outFilterMismatchNmax 999), minimum intron size of 20 bp, maximum intron size of 1,000,000 bp, minimum splice junction overhang of 8 bp for unannotated junctions (–alignSJoverhangMin 8), and minimum 1 bp overhang for annotated splice junctions (–alignSJDBoverhangMin 1). Gene annotations were obtained from GENCODE v44.

***Gene Quantification.*** Gene-level read counts were generated during alignment using STAR’s built-in quantification mode (–quantMode GeneCounts). Unstranded counts (column 2 of STAR’s ReadsPerGene.out.tab output) were extracted and assembled into a count matrix for all samples using Ensembl gene identifiers.

***Differential Expression Analysis.*** Differential expression analysis was performed using DESeq2. Genes with fewer than 10 total counts across all samples were filtered out prior to analysis. The standard DESeq2 workflow was applied with design formula ∼condition, comparing TES versus GFP (GFP set as the reference level). P-values were adjusted for multiple testing using the Benjamini-Hochberg method. Genes with adjusted p-value < 0.05 were considered significantly differentially expressed. Variance stabilizing transformation (VST) was applied for visualization purposes including principal component analysis (PCA) and heatmaps. Gene symbols were mapped from Ensembl IDs using the org.Hs.eg.db annotation package.

***Functional Enrichment Analysis.*** Gene Ontology (GO) enrichment analysis was performed separately for upregulated and downregulated genes using clusterProfiler. GO terms from Biological Process (BP), Molecular Function (MF), and Cellular Component (CC) ontologies were tested with Benjamini-Hochberg correction (p-value cutoff 0.05, q-value cutoff 0.2). Gene Set Enrichment Analysis (GSEA) was performed using clusterProfiler with genes ranked by the signed −log10(p-value) multiplied by the sign of the log2 fold change. Gene sets from MSigDB were tested including Hallmark gene sets, GO Biological Process, Reactome pathways, KEGG pathways, and Oncogenic signatures. GSEA parameters included minimum gene set size of 15, maximum gene set size of 500, and 10,000 permutations.

***Transcription Factor Binding Site Enrichment in DEG Promoters.*** To identify transcription factor binding motifs enriched at differentially expressed gene promoters, HOMER findMotifsGenome.pl was used. Promoter regions were defined as TSS ± 2,000 bp for all genes in the genome using GENCODE v44 annotations. Differentially expressed genes (adjusted p-value < 0.05) were split into upregulated (log2 fold change > 0) and downregulated (log2 fold change < 0) sets, and motif enrichment was performed separately for each set. Background regions consisted of promoters of all annotated genes. De novo motif discovery was performed for motif lengths of 8, 10, and 12 bp (-len 8,10,12), using exact BED region coordinates (-size given), the hg38 genome reference, and the top 25 motifs were reported.

### CUT&Tag analysis

***Quality Control and Preprocessing.*** Raw sequencing reads were assessed for quality using FastQC. Adapter sequences and low-quality bases were removed using Trim Galore in paired-end mode (–paired) with quality threshold of Phred 20, adapter detection stringency of 5 bp, minimum read length of 20 bp, and gzip-compressed output (–gzip). FastQC was run on the trimmed reads (–fastqc). CUT&Tag data generated in this study are accessible through the GEO database with the GEO series accession number GSE319889.

***Read Alignment.*** Trimmed reads were aligned to the human reference genome (GRCh38) using Bowtie2 with Cut&Tag-optimized parameters: very-sensitive alignment mode (–very-sensitive), proper pairs only (–no-mixed –no-discordant), insert size range of 10-700 bp (-I 10 -X 700), and Phred33 quality encoding.

***Filtering and Deduplication.*** Aligned reads underwent stringent quality filtering. Read groups were added using Picard AddOrReplaceReadGroups. PCR duplicates were identified and removed using Picard MarkDuplicates (REMOVE_DUPLICATES=true). Alignments were filtered using SAMtools for: properly paired reads (SAM flag 2), high mapping quality (MAPQ >= 30), and exclusion of unmapped reads, mate unmapped, secondary alignments, failed QC, and duplicates (-F 1804, corresponding to SAM flags 4, 8, 256, 512, and 1024). Analysis was restricted to canonical chromosomes (1-22, X, Y), excluding mitochondrial DNA and unplaced contigs.

***Peak Calling.*** Narrow peaks were called using MACS2 in paired-end mode using BAMPE format (-f BAMPE), which uses actual fragment coordinates from paired-end reads without model building or fragment extension. Additional parameters included: hg38 effective genome size (-g 2913022398), q-value threshold of 0.05 (-q 0.05), and retention of all reads (–keep-dup all). Mouse IgG was used as control for TES samples, and rabbit IgG for TEAD1 samples. Called peaks were filtered to retain only those with fold enrichment >= 2 (narrowPeak column 7). Peaks overlapping ENCODE blacklist regions (hg38-blacklist.v2.bed) were removed using bedtools intersect (-v). Consensus peaks were generated by calling peaks on combined replicates, and individual replicate peaks were called for reproducibility assessment.

***Coverage Track Generation.*** Normalized coverage tracks (BigWig format) were generated using deepTools bamCoverage at 10 bp resolution (–binSize 10) with read extension (–extendReads). Two normalization methods were applied: counts-per-million (CPM) and reads per genomic content (RPGC, –effectiveGenomeSize 2913022398) for visualization in genome browsers.

***Peak Annotation.*** Peaks were annotated to genomic features using ChIPseeker with the TxDb.Hsapiens.UCSC.hg38.knownGene annotation database. The promoter region was defined as −3,000 to +3,000 bp from transcription start sites. Gene symbols were obtained using the org.Hs.eg.db annotation package.

***Differential Binding Analysis.*** Differential binding analysis between TES and TEAD1 was performed using DiffBind with DESeq2 as the underlying statistical method. Consensus peak sets were generated from replicate overlap analysis, and default normalization was applied.

***Motif Analysis.*** *De novo* motif discovery and known motif enrichment analysis were performed on Cut&Tag peaks using HOMER findMotifsGenome.pl with the following parameters: 200 bp region size centered on peak summits (-size 200), repeat masking (-mask), and hg38 genome reference.

### Integration analysis

**Direct Target Identification.** Direct transcriptional targets of TES and TEAD1 were identified by integrating Cut&Tag binding data with RNA-seq differential expression data. Peaks were annotated using ChIPseeker, with promoter regions defined as −3,000 to +3,000 bp from the transcription start site. A gene was classified as a direct target if it met both criteria: (1) significant differential expression (adjusted p-value < 0.05) in RNA-seq comparing TES versus GFP, and (2) presence of a TES or TEAD1 binding peak in the promoter region (within 3 kb of the TSS, as defined by ChIPseeker annotation) or in a distal regulatory region within 50 kb of the TSS. Genes meeting only the expression criterion were classified as indirect targets, potentially regulated through secondary transcriptional effects.

***Functional Analysis of Target Genes.*** GO enrichment analysis was performed on direct target gene sets using clusterProfiler with Benjamini-Hochberg correction to identify enriched biological processes among genes directly bound and regulated by TES/TEAD1.

### Animals

Six-week-old male and female NSG mice (NOD.cg-Prkdc scid Il2rgtm1Wjl/SzJ) and C57BL/6j wild type mice were purchased from Jackson Laboratories. Upon arrival, mice were maintained at the San Raffaele Scientific Institutional Mouse Facility. Animals were group-housed in a 12 hours light/dark cycle and with ad libitum access to food and water. All the animal holding rooms were maintained to a controlled temperature of 25°C and to a relative humidity of 50% ± 5%. All experiments were performed in accordance with experimental protocols approved by local Institutional Animal Care and Use Committee. All efforts were made to keep animal usage to a minimum, in accordance to the achievement of the study aims.

### Human xenografts in mice

CSCs L0627 were used to generate human xenografts in mice. In particular, when preinfected cells were injected, CSCs L0627 were seeded in 6-multiwell plate and infected with 10 uL of LV expressing TES or GFP (1×10^9 IFU/mL) for each well. After 48h cells were collected and prepared for injections.

***Heterotopic xenografts.*** A total of 3×10^6 infected cells were resuspended in 100 µL of growth factor–reduced Matrigel (Corning). Using 1 ml syringes pre-cooled at −20°C, GFP-infected cells were subcutaneously injected into the left flank of NSG mice (NOD.cg-Prkdc scid Il2rgtm1Wjl/SzJ), whileTES-infected cells were subcutaneously injected into the right flank of the same animal. Mice were sacrificed one month after injection, in accordance with CSCs growth kinetics, and subcutaneous tumors were excised and fixed in 4% paraformaldehyde (PFA, Sigma-Aldrich) for 24 hours. Tumor samples were then measured, kept overnight in 30% Sucrose in PBS (Sigma-Aldrich), and embedded in optimal cutting temperature compound (O.C.T., VWR, 361603E) for cryopreservation. After flash frozing, serial coronal sections of 50 µm thickness were obtained using a cryostat (CM1850 UV, Leica). Subsequently, sections were processed for immunofluorescence analysis or mounted on gelatin-coated glass slides and processed for Nissl staining.

***Intracranial xenograft***. A total of 3×10^5 infected cells were resuspended in 3 µL of 1X PBS and then were unilaterally injected into the right striatum of 6-week-old NSG mice (male and female) at a controlled flow rate of 0.5 µL/min. Stereotaxic injections were performed using the following coordinates from the bregma: antero-posterior (AP) +0.5 mm, medio-lateral (ML) −1.8 mm, dorso-ventral (DV) −3.3 mm from the skull surface. Animals were maintained under isoflurane anesthesia throughout the stereotaxic surgical procedure. Mice were sacrificed at 2, 3 and 5 weeks from injection (always upon clinical assessment of animal welfare). Following deep anesthesia, mice were transcardially perfused with 4% PFA in PBS, then brains were removed from the skull and post-fixed overnight in 4% PFA. After fixation, brains were kept overnight in 30% sucrose in PBS and then embedded in O.C.T. for cryopreservation. Serial coronal sections of 50 µm thickness were obtained using a cryostat (CM1850 UV, Leica). Subsequently, sections were processed for immunofluorescence analysis or mounted on gelatin-coated glass slides and processed for Nissl staining.

***In vivo treatment.*** Orthotopic intracranial xenograft of naive CSCs L0627 were induced as described before. After 7 days from cell injection, the mice, randomly divided in two groups, were unilaterally injected at the same controlled flow rate and topological coordinates with 3 uL of LVs carrying GFP or TES (1×10^9 IFU/mL). Animals were maintained under isoflurane anesthesia throughout the stereotaxic surgical procedure. After 3 weeks from LV injections, mice were anesthetized and transcardially perfused with 4% PFA in PBS, then brains were removed from the skull and post-fixed overnight in 4% PFA. After fixation, brains were kept overnight in 30% sucrose in PBS and then embedded in O.C.T. for cryopreservation. Serial coronal sections of 50 µm thickness were obtained using a cryostat (CM1850 UV, Leica). Subsequently, sections were processed for immunofluorescence or mounted on gelatin-coated glass slides and processed for Nissl staining.

***Survival assessment.*** Orthotopic intracranial xenograft of naive CSCs L0627 were induced as described before. After 12 days from cell injection, all the mice were treated with 50mg/Kg/die of Temozolomide (TMZ, Sigma-Aldrich, T2577) for 3 days through oral gavage. After one day of wash-out from the treatment, the animals were randomly divided in two groups and were unilaterally injected into the striatum with 3 uL of LVs carrying GFP or TES (1×10^9 IFU/mL) at the same controlled flow rate and topological coordinates as described before. Animals were maintained under isoflurane anesthesia throughout the stereotaxic surgical procedure. Mice were monitored for survival analysis and euthanized upon clinical signs indicating compromised general welfare. For sacrifice, mice were anesthetized and transcardially perfused with 4% PFA in PBS, then brains were removed from the skull and post-fixed overnight in 4% PFA. After fixation, brains were kept overnight in 30% sucrose in PBS and then embedded in O.C.T. for cryopreservation. Serial coronal sections of 50 µm thickness were obtained using a cryostat (CM1850 UV, Leica). Subsequently, sections were processed for immunofluorescence analysis.

***Migration behaviour in intracranial xenograft.*** Orthotopic intracranial xenograft of TES or Mock infected CSCs L0627 were induced as described before. For this experiment, GFP-labelled CSCs cells were used for transplantation. Mice were sacrificed at 2 weeks from cells injection (always upon clinical assessment of animal welfare). Following deep anesthesia, mice were transcardially perfused with 4% PFA in PBS, then brains were removed from the skull and post-fixed overnight in 4% PFA. After fixation, brains were kept overnight in 30% sucrose in PBS and then embedded in O.C.T. for cryopreservation. Serial coronal sections of 50 µm thickness were obtained using a cryostat (CM1850 UV, Leica). Subsequently, sections were processed for immunofluorescence and immunohistochemistry analysis. To assess tumor cell migration from the primary tumor mass, regions of interest (ROIs) were manually delineated around the tumor core in each section. Concentric ROIs were then projected outward at 10 µm intervals from the primary tumor boundary (0 µm) up to 100 µm. Cells located within each 10 µm “ring” were counted to quantify the number of migrated cells at increasing distances from the tumor core. For each tumor, migration was quantified both as the absolute number of cells within each “ring”, and as the percentage of migrating cells at each distance relative to the total number of migrated cells (sum of all cells migrated from 0 to 100 µm).

No statistical methods were used to predetermine sample sizes. Mice were randomized before the procedures. Data collection and analyses were not performed blind to the conditions.

### Viral administration in wild-type mice

Unilateral hippocampal injections were performed in wild-type (WT) C57BL/6 mice of both sexes to assess the safety of TES in healthy brain tissue. Briefly, two injections per hippocampus (0.8 µL each; controlled flow rate of 0.5 µL/min) of lentiviral vectors expressing GFP or TES (1×10^9 IFU/mL) were carried out at the following stereotaxic coordinates: AP −2.8 mm, ML - 3 mm, DV −3.5 mm and −2.5 mm. Animals were maintained under isoflurane anesthesia throughout the stereotaxic surgical procedure. One month after the surgery, mice were sacrificed and transcardially perfused as previously described, and brains were collected and processed for molecular and histological analyses.

### Immunostaining

Cells were seeded onto glass coverslips (previously coated with Matrigel for CSC cultures to promote cell adhesion) and fixed on ice for 20 min in 4% paraformaldehyde (PFA, Sigma-Aldrich) in phosphate-buffered saline (PBS, Euroclone). After fixation, cells were washed with PBS and permeabilized for 1 hour in blocking solution containing 0.1% Triton X-100 (SigmaAldrich) and 10% donkey serum (Euroclone). Cells were then incubated overnight at 4 °C with primary antibodies diluted in blocking solution. The following day, cells were washed three times with PBS (5 min each) and incubated for 1 h at room temperature with secondary antibodies and Hoechst 33342 (ThermoFisher Scientific) diluted in blocking solution. Immunostaining of brain sections was performed on free-floating condition. Briefly, brain sections were permeabilized for 10 min with a solution of 10% methanol and 3% hydrogen peroxide (Sigma-Aldrich) in PBS and 20 min with 2% Triton X-100 in PBS. After three washes in PBS, slices were blocked for 1 h at room temperature in a solution containing 3% bovine serum albumin (BSA) and 0.3% Triton X-100 in PBS to saturate unspecific sites. Incubation with primary antibodies was performed at 4°C overnight in blocking solution containing 1% BSA and 0.1% Triton X-100 in PBS. The following day, appropriate secondary antibodies were applied to sections for 1 h at room temperature in the same blocking solution supplemented with Hoechst 33342. Both cell coverslips and brain sections were subsequently washed and mounted using Fluorescent Mounting Medium (Dako, S3023). Images were acquired using a Nikon DS-Qi2 epifluorescence microscope or Leica SP8 and Mavig RS-G4 confocal microscopes, and analyzed with ImageJ/Fiji software. For the immunohistochemical analysis, following overnight incubation with the primary antibody, brain slices were incubated with an appropriate biotinylated secondary antibody. Subsequently, sections were treated with an avidin-biotin-enzyme complex and immunoreactivity was revealed using 3,3’-diaminobenzidine solution (DAB, Vector Laboratories) as chromogen, according to the manufacturer’s instructions. Finally, slices were mounted for imaging using the Eukitt® mounting medium (Sigma-Aldrich). Images were acquired using Zeiss Axio Imager M2m microscope and analyzed with ImageJ/Fiji software. The primary antibodies used in this study were as follows: anti-V5 (1:500; mouse; Thermo Fisher Scientific, R96025), anti-GFP (1:1000; chicken; Thermo Fisher Scientific, A10262), anti-KI67 (1:500; clone SP6, rabbit; Immunological Sciences, MAB-90948), anti-MAP2 (1:1000; chicken; Abcam, ab92434), anti–phospho-histone H3 (PH3, 1:200; Ser10, rabbit; Sigma-Aldrich, 06-570), anti-cleaved caspase 3 (CC3, 1:200; Asp175, rabbit; Cell Signaling Technology, 9661), anti-human nuclear antigen (1:100; rabbit; Thermo Fisher Scientific, RBM5-346-P1). A complete list of antibodies used in this study is provided in Table S8. For propidium iodide staining, cell cultures were live-stained prior to fixation with Propidium Iodide (1µg/µL, ThermoFisher Scientific, P3566) and RNase A (100 µg/mL, Sigma-Aldrich) solution in PBS at 37 °C for 20 min in the dark. Cells were subsequently processed for immunofluorescence as described above. For immunofluorescence analysis on cultured cells, replicates shown in bar graphs represent the number of coverslips from independent platings used for each quantification. For tissue immunofluorescence, replicates correspond to the number of animals analyzed.

### Nissl staining

Brain sections were rinsed in distilled water for 1 min and subsequently stained with 0,1% Cresyl Violet solution (Sigma-Aldrich, C5042) preheated at 50°C for 7 min. Sections were then rinsed in distilled water for 3 min and dehydrated through an ascending alcoholic scale (70% - 100% ethanol) for 1 min each. Finally, brain slices were cleared in xylene for 2 hours and mounted for imaging using the Eukitt® mounting medium (Sigma-Aldrich).

### Primary mouse neuronal cultures

Primary cultures from mouse embryonic cortices were prepared from E17.5 (embryonic day 17.5) CD1 WT mice. Briefly, following sacrifice, brains were isolated and the cerebral cortices were dissected by removing the meninges and separating the hemispheres along the dorsal midline. Cortices were then enzymatically digested with 0.025% trypsin (GIBCO) in Hank’s Balanced Salt Solution (HBSS, Euroclone) for 20 min at 37°C. Trypsin was then removed and cortices were washed with plating medium (Neurobasal A medium supplemented with 1x B-27, 3.3 mM glucose, 2mM glutamine and 1% Pen/Strept) and mechanically dissociated using a P1000 pipette to obtain a homogeneous cell suspension. Cells were plated onto poly-L-lysine (PLL, 0.1 mg/ml) coated glass coverslips. After 24h, primary cultures were transduced with 2 uL of lentiviruses expressing TES or GFP, and 48 h after infection a subset of coverslips was fixed as previously described to assess transduction efficiency by immunocytochemical analyses. Additional coverslips were fixed at 21 days post-infection to evaluate neuronal survival and morphological features over time.

### In silico modeling

To model TES fusion protein, the full-length amino acid sequences was entirely submitted to I-TASSER^29,30^, a 3D structure predictor, based on the homology modeling algorithm. From the software output, the top five models were kept and ranked by I-TASSER score (C-score) and root mean square deviation of atomic positions, that allow to know the general folding rate of a protein structure, for further analysis (Table S1). To assess whether the ability to bind DNA is maintained, a blind docking simulation analysis was performed using the predicted TES and double helix DNA strand within Haddock software^31^. The DNA used as ligand was extracted from the ProteinData Bank file 5NNX (the ternary complex of the DNA binding domains of TEAD1 with a 18-bp oligonucleotide). The top 10 poses ranked by 𝚫G of binding (kcal/mol) were reported (Table S1).

### Statistical analysis

Statistical analysis was performed using GraphPad Prism (GraphPad software, San Diego, CA, USA; version 9.0). Values are expressed as mean ± SEM as indicated and significance was set at *p* < 0.05. Statistical differences were determined by analysis of variance (ANOVA test), by Survival or by Student’s two-sided Unpaired t test followed, when significant, by an appropriate post hoc test. In all reported statistical analyses, differences were labeled as non-significant for p > 0.05, significant (*) for p < 0.05, very significant (**) for p < 0.01 and extremely significant (***) for p < 0.001 and (****) for p < 0.0001. n ≥ 3 indicates the number of repeats for each individual experiment.

## Supplementary Table Legends

**Table S1. TES *in silico* model**

Sheet 1: I-TASSER output for the top five models.

Sheet 2: parameters of the docking obtained with Haddock for TES model.

**Table S2. RNA-sequencing in SNB19 cells**

List of DEGs (both adjusted p value < 0.05 and |log2FoldChange|>1).

**Table S3. Statistically enriched categories**

Statistically significant Gene Ontology (BP) categories and GSEA sets (Hallmark, KEGG, Oncogenic, Reactome).

**Table S4. YAP/TAZ target genes**

Manually curated targets of YAP/TAZ experimentally validated in literature.

**Table S5. CUT&Run peaks**

Sheet 1: Table S5a, TES specific peaks. Sheet 2: Table S5b, TEAD1 specific peaks.

**Table S6. Cell lines used in this work**

List of cell lines used within this work. **Table S7. Antibodies used in this work** List of antibodies used within this work.

**Table S8. Oligonucleotides used in this work**

List of primers used within this work.

## Supporting information

Table S1. TES in silico model

Table S2. RNA-sequencing in SNB19 cells

Table S3. Statistically enriched categories

Table S4. YAP/TAZ target genes

Table S5. CUT&Run peaks

Table S6. Cell lines used in this work

Table S7. Antibodies used in this work

Table S8. Oligonucleotides used in this work

## ACKNOWLEDGMENTS

We are thankful to C. Mussolino for providing plasmids and to R. Galli for sharing patient-derived CSCs and advice. We thank the members of the Broccoli and Sessa Labs for helpful discussions.

Funding: this work was supported by Repron Therapeutics to V.Br. and A.S., and by AIRC foundation (IG2025 - ID 32238) to V.Br. and (MFAG2021 - ID 26017) to A.S.

## AUTHOR CONTRIBUTIONS

V.Br. and A.S. conceived the study and designed the experiments. E.B, S.F., A.C., V.Be., and F.B. performed and analyzed all the experiments with the help of S.G.G., M.L., and E.V. Bioinformatics analysis was performed by M.K., and E.B.. G.C. and F.U. provided reagents and advice. V.Br. and A.S. wrote the manuscript and provided financial support. All authors revised the manuscript.

---

**Figure S1.**
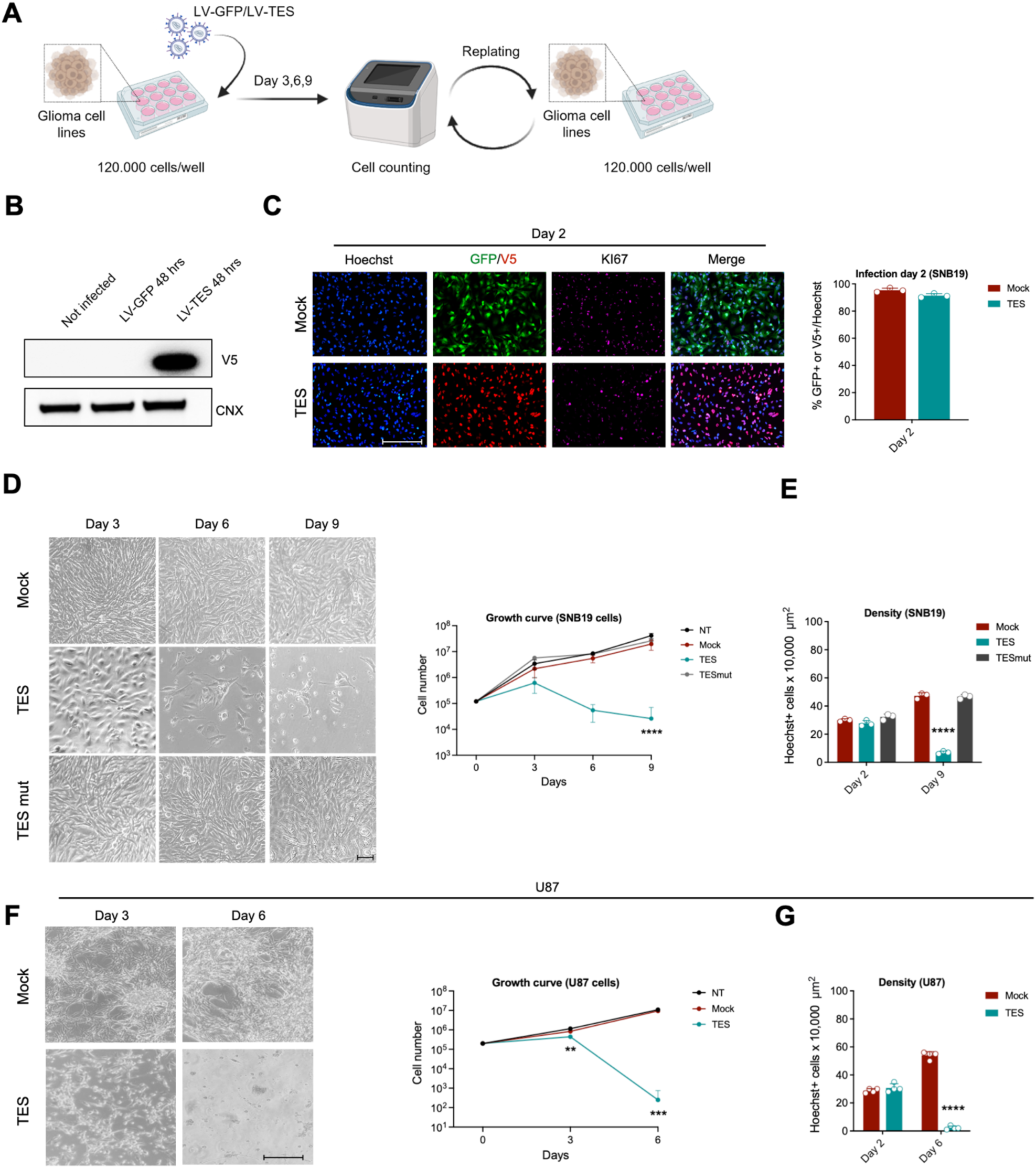
TES *in vitro* efficacy on human glioma cell lines. (A) Experimental scheme of the infection protocol used in human glioma cell lines (SNB19 and U87). (B) Western blot for V5 and calnexin (CNX) (as loading control) in SNB19 cells not infected or infected with lentivirus carrying GFP or TES, confirming the presence of TES protein expression in cells infected. Proteins were extracted 48 hours after infection. (C) Immunocytochemistry for Ki67 mitotic marker and for GFP or V5 tag, counterstained with Hoechst, in SNB19 cells performed 48 hours after GFP or TES lentiviral infection. Quantification is represented as percentage of GFP+ or V5+ cells on the total number of Hoechst+ nuclei (means ± SEM) in order to verify infection efficiency. Mock vs TES *P*=0.1201. Statistical analysis was performed using paired *t* test. *n*=3. Scale bar: 250 μm. (D) Representative bright-field images and growth curve of SNB19 cell line infected with GFP (mock, in red), TES (in blue), TESmut (in grey), or not treated cells (NT, in black), indicating strong capability of TES in inhibiting tumor cell growth and proliferation, and the inactivity of TESmut probably because of its inability to bind DNA on TEAD1 target sites. Growth curve: Mock vs TES Day 3 *P*=0.8868; Mock vs TESmut Day 3 *P*=0.4252; Mock vs TES Day 6 *P*=0.1262; Mock vs TESmut Day 6 *P*=0.6517; Mock vs TES Day 9 \*\*\*\**P*<0.0001; Mock vs TESmut Day 9 *P*=0.0506. Data are expressed as means ± SEM. Statistical analysis was performed using two-way ANOVA followed by Dunnett’s multiple comparisons test. *n* = 3. Scale bar: 50 μm. (E) Quantification of Hoechst+ cells x 10,000 μm^2^ (Area=1 cm^2^) at 2 and 9 days post infection with GFP (mock, in red), TES (in blue), or TESmut (in grey) in SNB19 cell line. Hoechst+ nuclei were counted within a fixed area of 100 μm ×100 μm to obtain the relative cell density: Mock vs TES Day 2 *P*=0.1498; Mock vs TESmut Day 2 *P*=0.1498; Mock vs TES Day 9 \*\*\*\**P*<0.0001; Mock vs TESmut Day 9 *P*=0.7670. Data are expressed as means ± SEM. Statistical analysis was performed using two-way ANOVA followed by Dunnett’s multiple comparisons test. *n* = 3. (F) Representative bright-field images and growth curve of U87 cell line infected with GFP (mock, in red), TES (in blue), or not treated cells (NT, in black), indicating strong capability of TES in inhibiting tumor cell growth and proliferation. Growth curve: Mock vs TES Day 3 \*\**P*=0.0011; Mock vs TES Day 6 \*\*\**P*=0.0006. Data are expressed as means ± SEM. Statistical analysis was performed using two-way ANOVA followed by Dunnett’s multiple comparisons test. *n* = 4. Scale bar: 100 μm. (G) Quantification of Hoechst+ cells x 10,000 μm^2^ (Area=1 cm^2^) at 2 and 6 days post infection with GFP (mock, in red) or TES (in blue) in U87 cell line. Hoechst+ nuclei were counted within a fixed area of 100 μm ×100 μm to obtain the relative cell density: Mock vs TES Day 2 *P*=0.4084; Mock vs TES Day 6 \*\*\*\**P*<0.0001. Data are expressed as means ± SEM. Statistical analysis was performed using two-way ANOVA followed by Sidak’s multiple comparisons test. *n* = 4.

**Figure S2.**
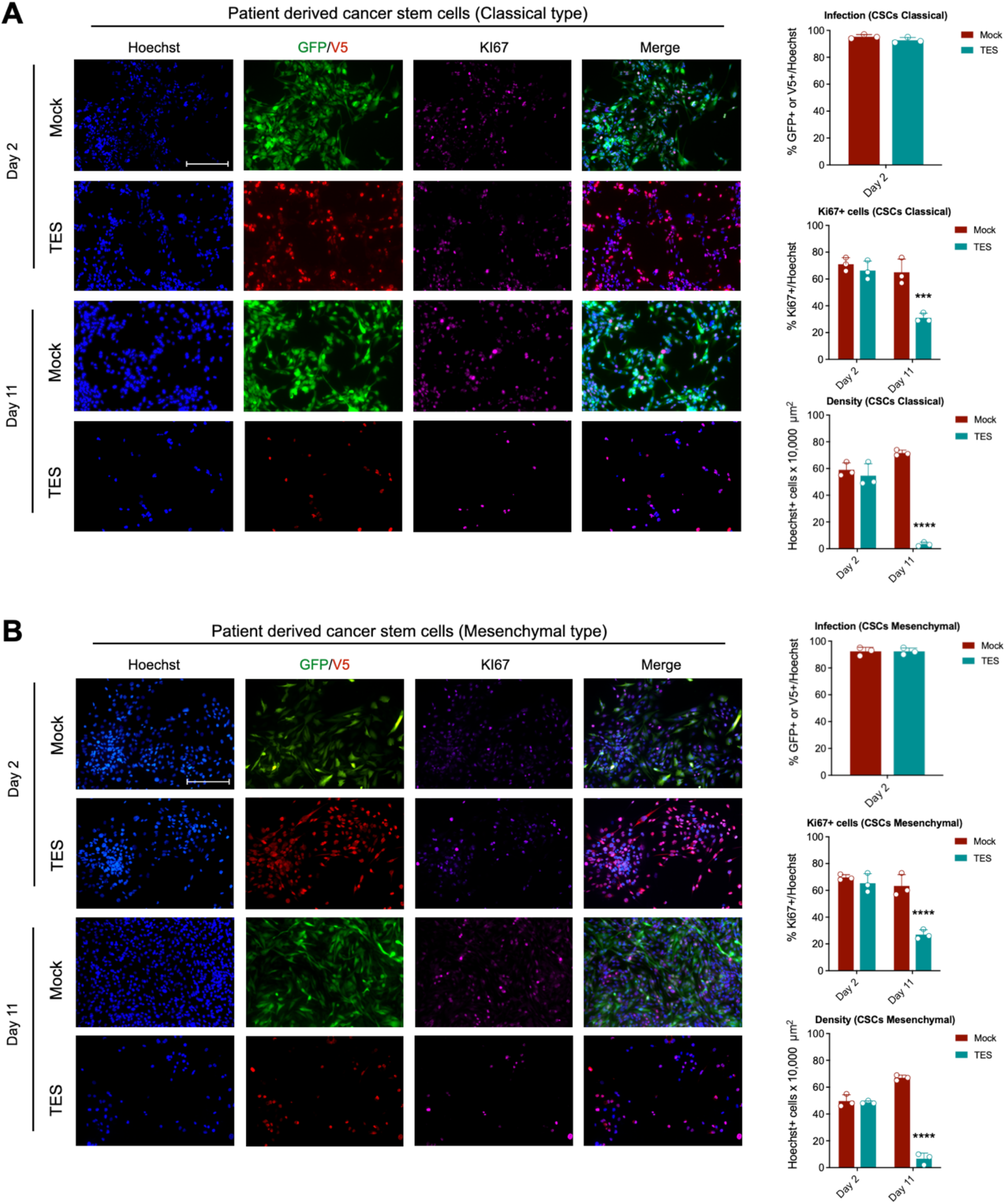
TES infection efficiency in patient-derived CSCs. (A) Immunocytochemistry for Ki67 mitotic marker and for GFP or V5 tag, counterstained with Hoechst, of patient-derived CSCs of classical GBM subtype performed at 2 *(upper)* and 11 *(lower)* days after GFP or TES lentiviral infection. Quantification is represented as percentage of GFP+ or V5+ cells on the total number of Hoechst+ nuclei (means ± SEM) in order to verify infection efficiency after 2 days: Mock vs TES *P*=0.1481. Statistical analysis was performed using unpaired *t* test. *n*=3. Quantification is represented as percentage of Ki67+ cells on the total number of Hoechst+ nuclei (means ± SEM), and indicates the ability of TES to reduce the number of proliferating cells: Mock vs TES Day 2 *P*=0.4243; Mock vs TES Day 11 \*\*\**P*=0.00028. Statistical analysis was performed using multiple *t* test. *n*=3. Quantification of Hoechst+ cells x 10,000 μm^2^ (Area=1 cm^2^) at 2 and 11 days post infection with GFP (mock, in red) or TES (in blue). Hoechst+ nuclei were counted within a fixed area of 100 μm ×100 μm to obtain the relative cell density: Mock vs TES Day 2 *P*=0.5791; Mock vs TES Day 11 \*\*\*\**P*<0.0001. Data are expressed as means ± SEM. Statistical analysis was performed using two-way ANOVA followed by Sidak’s multiple comparisons test. *n* = 3. Scale bar: 250 μm. (B) Immunocytochemistry for Ki67 mitotic marker and for GFP or V5 tag, counterstained with Hoechst, of patient-derived CSCs of mesenchymal GBM subtype performed at 2 *(upper)* and 11 *(lower)* days after GFP or TES lentiviral infection. Quantification is represented as percentage of GFP+ or V5+ cells on the total number of Hoechst+ nuclei (means ± SEM) in order to verify infection efficiency after 2 days: Mock vs TES *P*>0.9999. Statistical analysis was performed using unpaired *t* test. *n*=3. Quantification is represented as percentage of Ki67+ cells on the total number of Hoechst+ nuclei (means ± SEM), and indicates the ability of TES to reduce the number of proliferating cells: Mock vs TES Day 2 *P*=0.3959; Mock vs TES Day 11 \*\*\*\**P*<0.0001. Statistical analysis was performed using multiple *t* test. *n*=3. Quantification of Hoechst+ cells x 10,000 μm^2^ (Area=1 cm^2^) at 2 and 11 days post infection with GFP (mock, in red) or TES (in blue). Hoechst+ nuclei were counted within a fixed area of 100 μm ×100 μm to obtain the relative cell density: Mock vs TES Day 2 *P*=0.9241; Mock vs TES Day 11 \*\*\*\**P*<0.0001. Data are expressed as means ± SEM. Statistical analysis was performed using two-way ANOVA followed by Sidak’s multiple comparisons test. *n* = 3. Scale bar: 250 μm.

**Figure S3.**
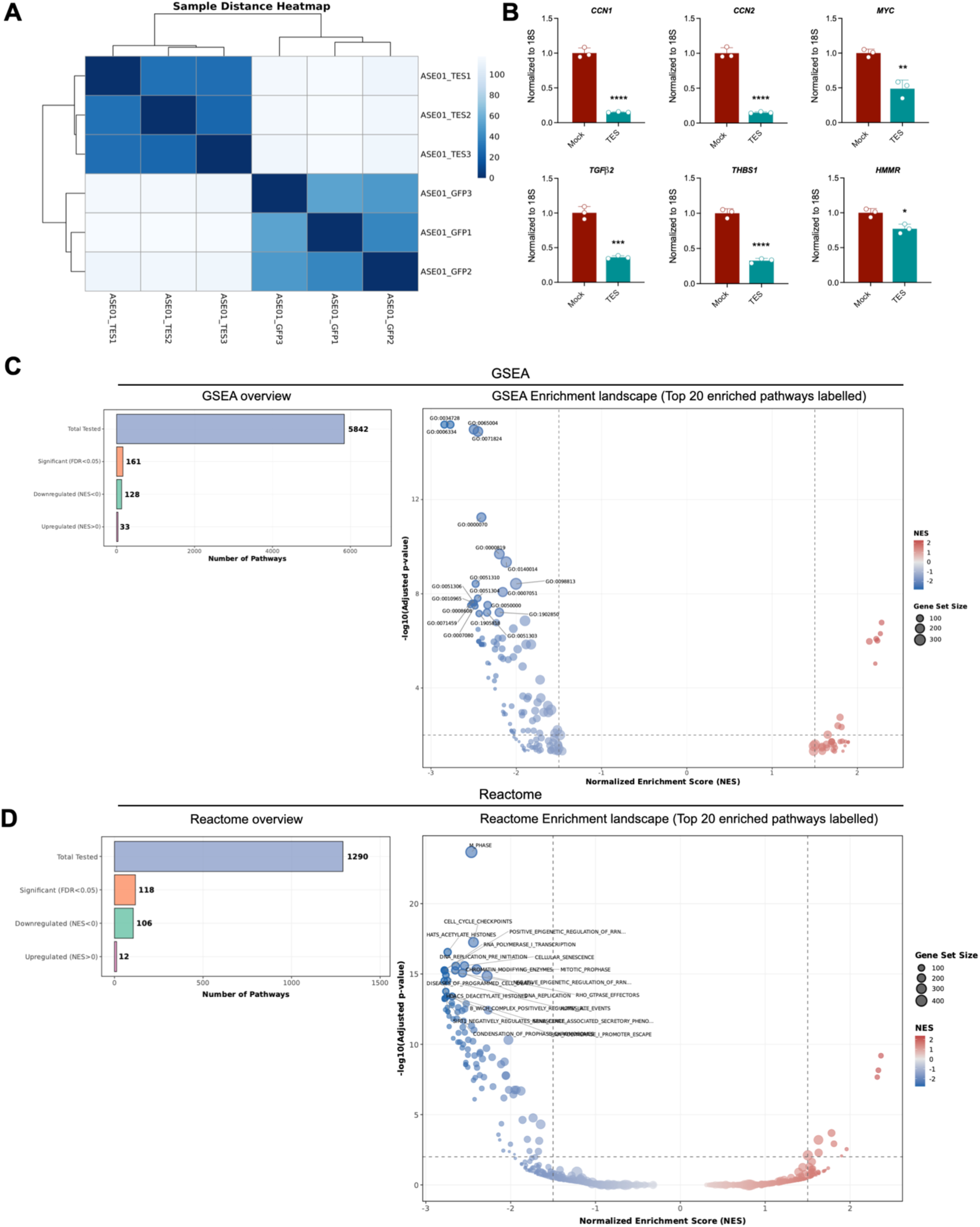
TES transcriptional activity in SNB19 cells. (A) Sample-to-sample distance heatmap of RNA-seq samples based on global gene expression profiles, highlighting clustering according to experimental conditions. (B) RT-qPCR on TEAD1 putative target genes (*CCN1, CCN2, MYC, TGFβ2*) and epithelial-to-mesenchymal transition related genes (*THBS1, HMMR*) at 2 days after GFP (mock) or TES lentiviral infection, confirming that TES treatment strongly downregulates their RNA expression. *CCN1*: \*\*\*\**P*<0.0001*; CCN2*: \*\*\*\**P*<0.0001*; MYC*: \*\**P*=0.0028*; TGFβ2*: \*\*\**P*=0.0003*; THBS1*: \*\*\*\**P*<0.0001; *HMMR*: \**P*=0.0108. Data are expressed as means ± SEM. Statistical analysis was performed using unpaired *t* test. *n* = 3. (C) GSEA analysis indicates that downregulated genes show largely more numerous significantly enriched terms, whereas genes upregulated show limited functional enrichment. (D) GSEA analysis for Reactome pathway identifies several significantly downregulated GO terms and gene sets associated with cell-cycle regulation.

**Figure S4.**
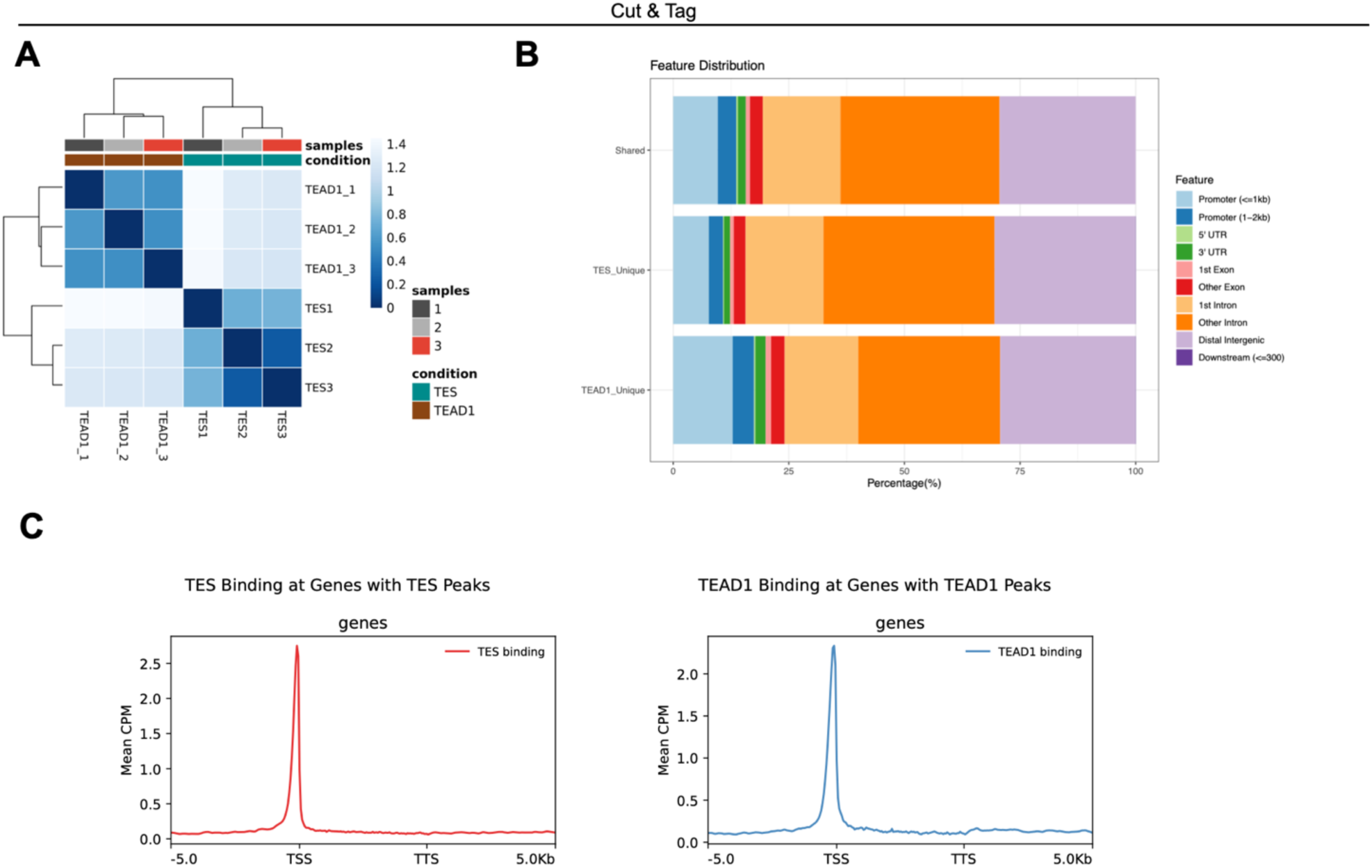
TES genome-wide occupancy. (A) Sample-to-sample distance heatmap of CUT&Tag samples based on genome-wide chromatin signal profiles, highlighting clustering according to experimental conditions. (B) Bar plot showing the distribution of TES only, TEAD1 only or TES/TEAD1 shared peaks across genomic features. All the three peak categories reported display similar genomic distributions. (C) Density plots showing the mean of TES or TEAD1 binding signal within ±5 kb around peak centers, normalized as counts per million (CPM).

**Figure S5.**
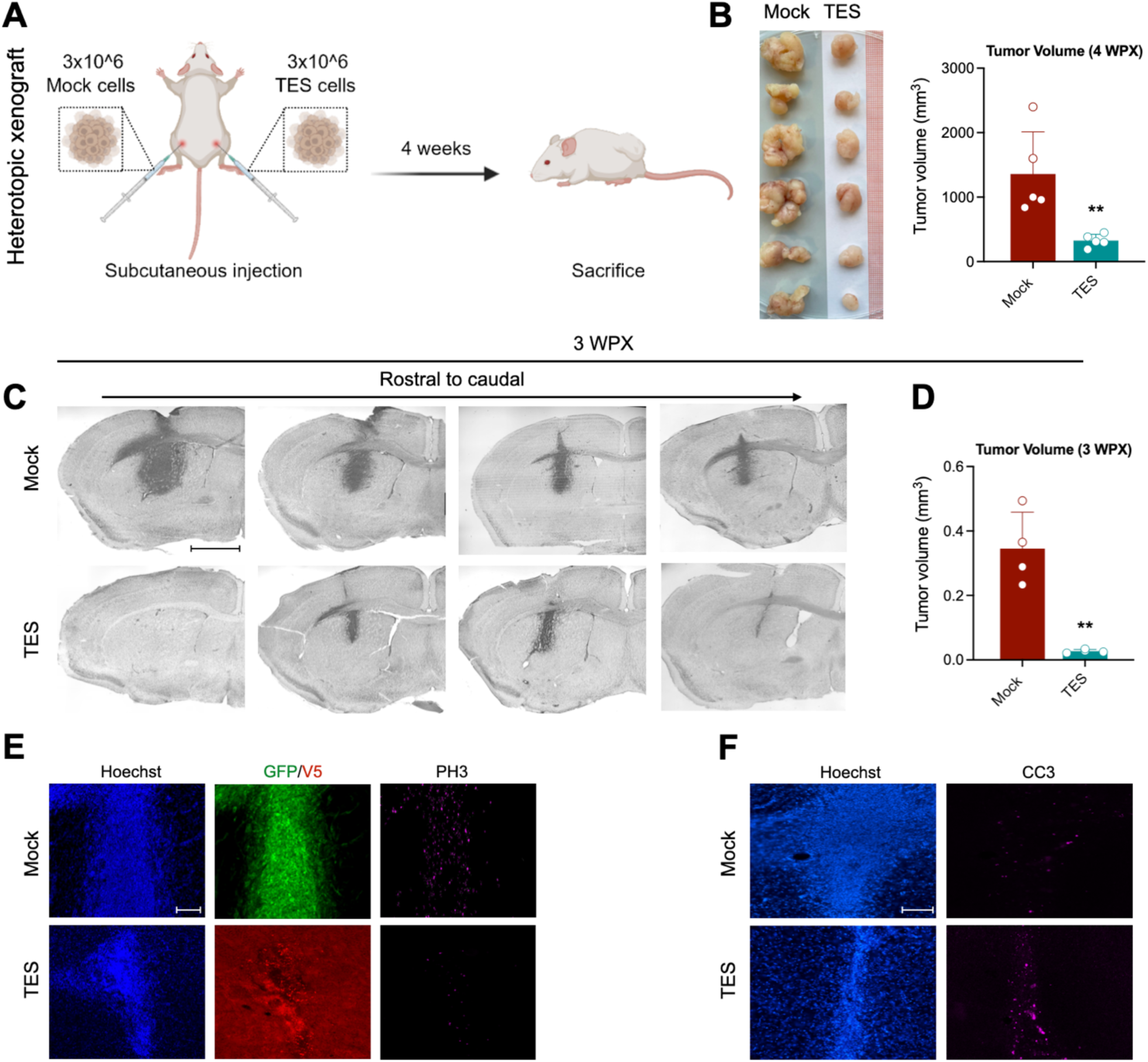
TES antitumor activity in mouse xenograft models. (A) Experimental scheme illustrating the generation of heterotopic xenografts by subcutaneous injections of Mock (left flank) or TES (right flank) pre-infected CSCs in immunodeficient NSG mice. (B) Photographs of the resulting tumors 4 weeks after subcutaneous injections: xenografts derived from TES-expressing cells are consistently smaller than those generated from GFP-expressing controls, indicating that TES expression attenuates tumor growth. Quantification of the tumor volume: Mock vs TES \*\**P*=0.0080. Data are expressed as means ± SEM. Statistical analysis was performed using unpaired *t* test. *n* = 5 animals per group. WPX, weeks post xenotransplantation. (C) Nissl staining of representative sections of the orthotopic xenograft model represented in Fig. 4A, 3 weeks after pre-infected CSCs transplantation, showing strong ability of TES in inhibiting tumor growth compared to Mock. Scale bar: 400 μm. (D) Quantification of tumor volume: Mock vs TES \*\**P*=0.0013. Data are expressed as means ± SEM. Statistical analysis was performed using unpaired *t* test. *n* = 4 animals per group. WPX, weeks post xenotransplantation. (E) Immunohistochemistry on representative coronal brain sections for GFP (Mock tag), V5 (TES tag) and PH3 (mitotic marker) counterstained with Hoechst in animals injected with Mock or TES pre-infected CSCs at 3WPX. Scale bar: 100 μm. (F) Immunohistochemistry on representative coronal brain sections for cleaved-caspase 3 (CC3, cell death marker) counterstained with Hoechst in same animals injected with Mock or TES pre-infected CSCs at 3WPX. Scale bar: 100 μm.

**Figure S6.**
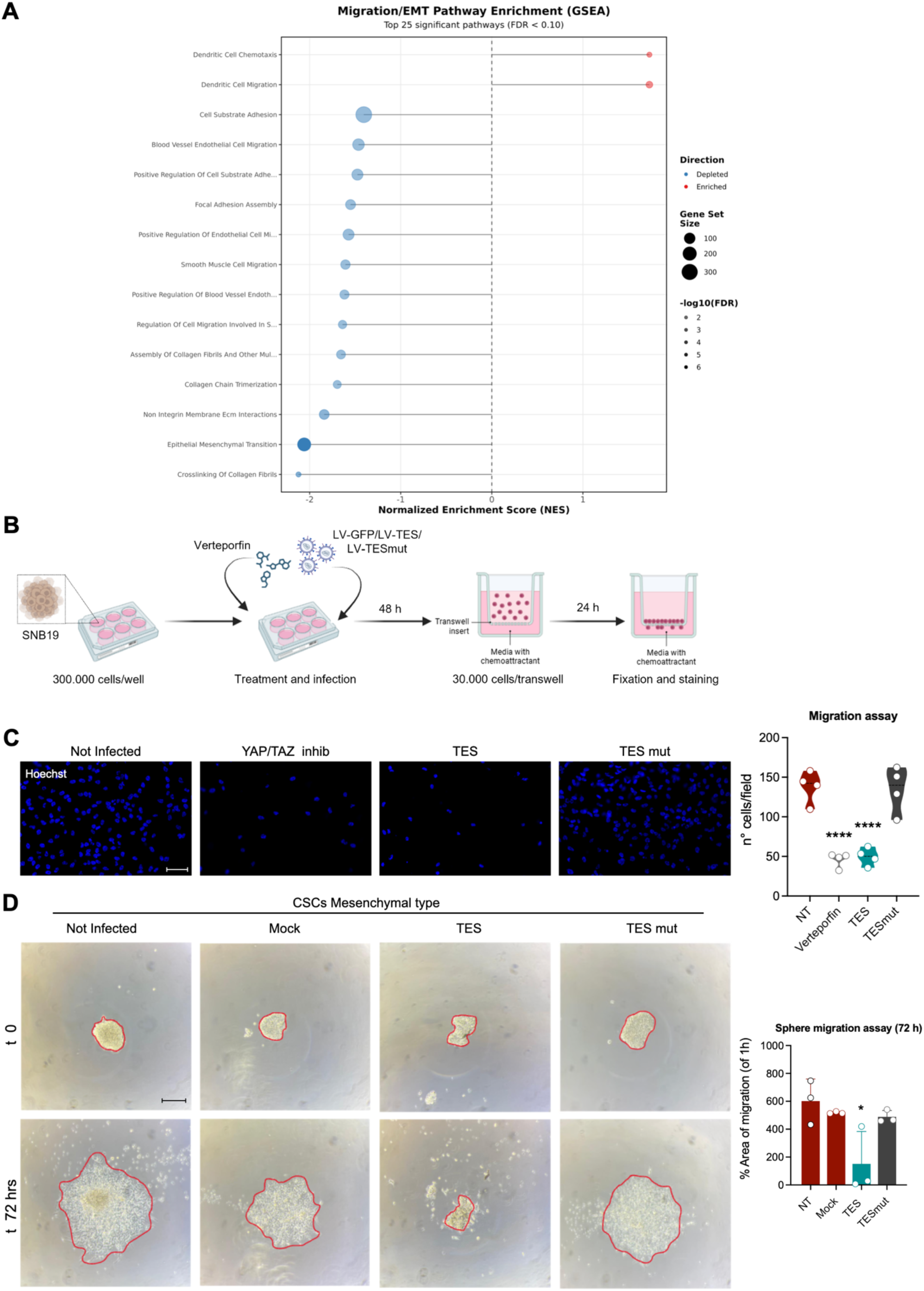
TES impairs cell migration. (A) GSEA analysis for migration/EMT pathway identifies several significantly downregulated GO terms and gene sets associated with migration regulation and processes. (B) Experimental scheme of the treatment protocol used in SNB19 cell line to perform transwell migration assay. (C) Hoechst staining of SNB19 cells not infected (NT), treated with Verteporfin (0.2 μM), infected with LV-TES and with LV-TESmut, at 24 hours after transwell plating, revealing TES capability of inhibiting cells migration, similarly to YAP/TAZ inhibitor verteporfin treatment. On the right, quantification of migrating cells to the bottom of the transwell at 24 hours: NT vs Verteporfin \*\*\*\**P*<0.0001; NT vs TES \*\*\*\**P*<0.0001; NT vs TESmut *P*=0.9874. Data are expressed as means ± SEM. Statistical analysis was performed using ordinary one-way ANOVA followed by Dunnett’s multiple comparisons test. *n* = 4. Scale bar: 100 μm. (D) Spheres diffusion assay shows decreased area of confluent cell migration (dispersion) at 72 hours in CSCs of mesenchymal subtype infected with LV-TES compared to Mock condition. Diffusion area is reported as percentage of area of migration normalized to the area of the sphere at t0 (t0 = 1 hour after spheres plating on PLL+laminin coating): Mock vs TES \**P*=0.0398; Mock vs TESmut *P*=0.8056. Data are expressed as means ± SEM. Statistical analysis was performed using ordinary one-way ANOVA followed by Holm-Sidak’s multiple comparisons test. *n* = 3. On the left, migration area of confluent cells is marked by red dash line in representative spheres. Scale bar: 200 μm.

